# MicroExonator enables systematic discovery and quantification of microexons across mouse embryonic development

**DOI:** 10.1101/2020.02.12.945683

**Authors:** Guillermo E. Parada, Roberto Munita, Ilias Georgakopoulos-Soares, Hugo Fernandez, Emmanouil Metzakopian, Maria Estela Andres, Eric A. Miska, Martin Hemberg

## Abstract

Microexons, exons that are ≤30 nucleotides, were shown to play key roles in neuronal development, but are difficult to detect and quantify using standard RNA-Seq alignment tools. Here, we present MicroExonator, a novel pipeline for reproducible *de novo* discovery and quantification of microexons. We processed 289 RNA-seq datasets from eighteen mouse tissues corresponding to nine embryonic and postnatal stages, providing the most comprehensive survey of microexons available for mouse. We detected 2,984 microexons, 332 of which are differentially spliced throughout mouse embryonic brain development, including 29 that are not present in mouse transcript annotation databases. Unsupervised clustering of microexons alone segregates brain tissues by developmental time and further analysis suggest a key function for microexon inclusion in axon growth and synapse formation. Finally, we analysed single-cell RNA-seq data from the mouse visual cortex and we report differential inclusion between neuronal subpopulations, suggesting that some microexons could be cell-type specific.

## Introduction

In eukaryotes, mRNA processing is a key regulatory step of gene expression (Hocine et al., 2010). Alternative splicing is arguably one of the most important processes affecting the vast majority of transcripts in higher eukaryotes (Licatalosi and Darnell, 2010). Consequently, alternative splicing impinges directly onto numerous biological processes such as cell cycle, cell differentiation, development, sex, circadian rhythm, response to environmental change, pathogen exposure and disease (Bell et al., 1988; Irimia and Blencowe, 2012; Kalsotra and Cooper, 2011; Venables et al., 2012). High-throughput RNA sequencing (RNA-seq) coupled with efficient computational methods have facilitated annotation of low abundance, tissue-specific transcripts and thus revolutionised our understanding of alternative and non-canonical splicing events (Sibley et al., 2016).

In vertebrates, dramatic changes in alternative splicing control neurogenesis, neuronal migration, synaptogenesis and synaptic function (Vuong et al., 2016). In particular, it was shown that short exons tend to be included more frequently in the central nervous system (Barash et al., 2010; Coelho and Smith, 2014). Recently, it was also shown that extremely short exons, known as microexons, herein defined as exons ≤30 nucleotides, are the most highly conserved component of neuronal alternative splicing during development (Irimia et al., 2014). Importantly, microexon inclusion has been proposed to have a key regulatory role during brain development, having an influence over neurite outgrowth, cortical layering and axon guidance (Quesnel-Vallières et al., 2015). However, the quantification of microexons using RNA-seq remains challenging due to their incomplete annotation (Ustianenko et al., 2017).

The first algorithms for genome-wide microexon discovery were based on EST/cDNA misalignment corrections, and discovered 170 microexons (Volfovsky et al., 2003; Wu and Watanabe, 2005). *De novo* discovery of microexon insertions by aligning short segments of mRNA using standard software is difficult because most algorithms require a perfectly matching seed sequence that often cannot fit within a single microexon. Detection can be improved by reducing the size of alignment seeds, as was done for Olego which enabled the identification of 630 novel microexons 9-27 nucleotides (nt) in mouse (Wu et al., 2013). Another strategy for increasing the sensitivity of microexon discovery is to directly map RNA-seq reads to libraries of annotated splice junctions (Irimia et al., 2014; Li et al., 2015), but the bioinformatic pipelines used in these seminal studies have not been released to the public domain. Today, VAST-TOOLS is the most widely used tool for microexon quantification from RNA-seq data (Tapial et al., 2017). However, a significant restriction of VAST-TOOLS is that it can only identify microexons that are annotated in VastDB (Tapial et al., 2017), which is only available for a limited number of species.

Here, we introduce MicroExonator, a computational workflow for discovery and quantification of microexons using RNA-seq data. MicroExonator employs a two-step procedure whereby it first carries out a *de novo* search for unannotated microexons and subsequently quantifies both new and previously annotated microexons (Fig 1). Using simulations we show that MicroExonator outperforms other available tools, both in terms of sensitivity and specificity. We then analyze mouse embryonic development RNA-seq datasets, and we identify a total of 2,984 microexons, 37% of these are not previously annotated in GENCODE (Harrow et al., 2006) mouse transcript models. We focus our analysis on 326 microexons that change during neuronal development, and 18% of which are not present in VastDB. Our analysis shows a pattern of orchestrated microexon inclusion during brain development as evidenced by the high degree of connectivity of the protein-protein interaction network encompassing the genes that contain microexons. We also directly demonstrate the high degree of conservation of microexons by analysing 23 zebrafish brain RNA-seq samples where we detect 348 zebrafish microexons that were conserved in mouse, including 54 that were not annotated in the Ensembl gene annotations. Finally, we apply MicroExonator to single-cell RNA-seq data, and we demonstrate that some microexons are not only tissue-specific, but also cell-type specific.

**Figure 1.**
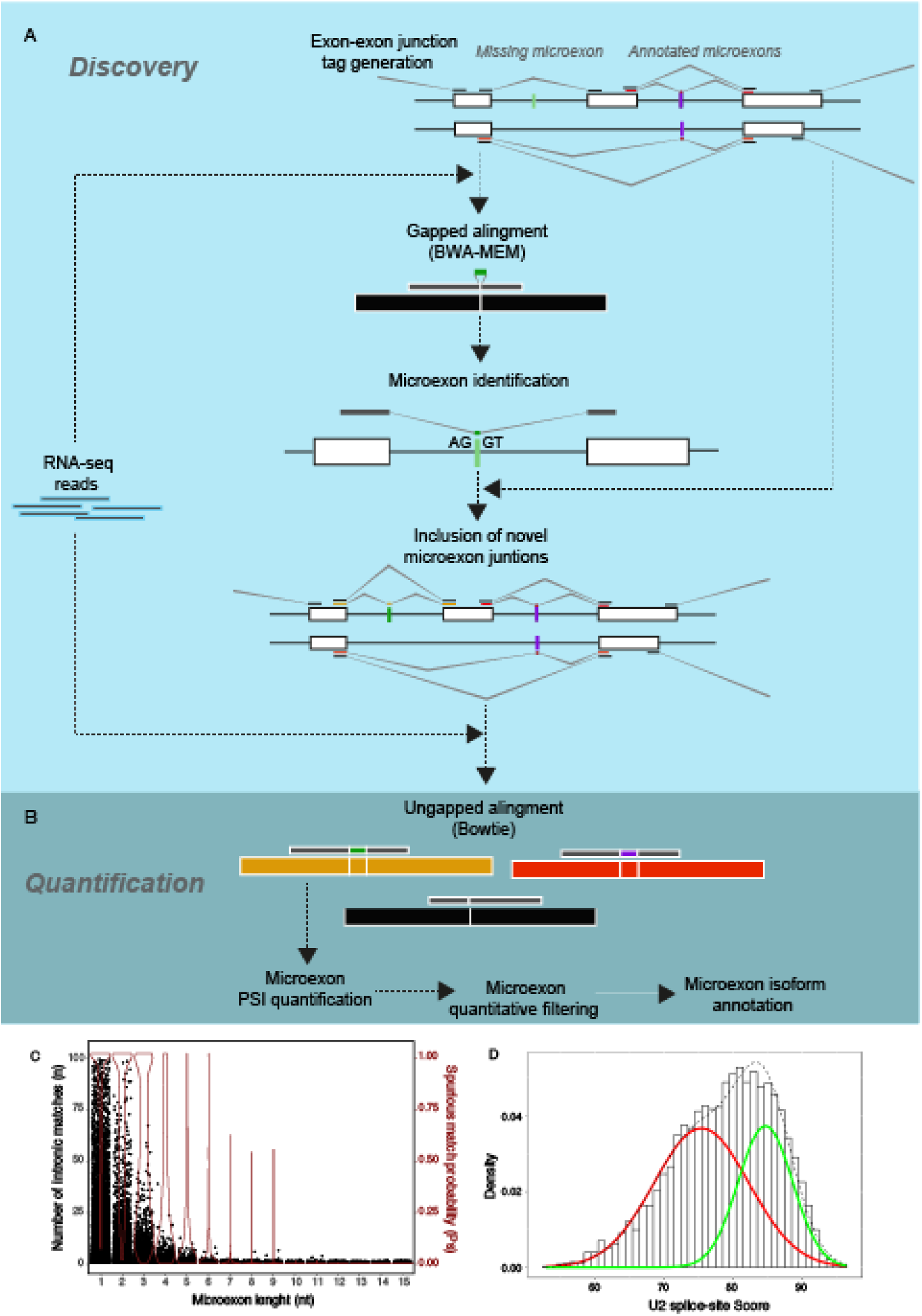
Overview of the MicroExonator workflow. A) To discover unannotated microexons, RNA-seq reads are aligned with BWA-MEM to the annotated splice junctions. The resulting alignments are post-processed to discover novel microexons flanked by canonical U2-type GT-AG splice sites. B) Both putative novel and annotated microexons are quantified and filtered to produce a final list of microexons into transcript models which can be used for downstream analysis. C) Number of intronic matches and distribution of spurious match probability across microexon lengths. D) A two-component Gaussian mixture is used to fit the U2 consensus splicing score distribution.

## Design

Microexons are a highly conserved and dynamically regulated set of cassette exons, and play an important role in neuronal development. However, microexons are often overlooked due to the difficulty of detecting them using standard RNA-seq aligners. Since MicroExonator is easy to incorporate into a RNA-seq processing workflow, it readily facilitates the investigation of microexons in transcriptome studies. Today, large volumes of RNA-seq data is publicly available, and it is not uncommon for researchers to analyse tens or hundreds of experiments in a single study. Such large scale analyses require an improved workflow, and when one relies on software packages written and maintained by others, reproducibility is an issue since different versions of software may not yield the same output. MicroExonator is based on state-of-the-art scientific software design principles (Grüning et al., 2018a), and it is implemented using SnakeMake (Köster and Rahmann, 2012), a widely used, high-performance workflow management systems (Larsonneur et al., 2018).

MicroExonator can be configured to download and process any number of RNA-seq samples that can be found locally or deposited on public archives, such as Short Read Archive, European Nucleotide Archive or ENCODE. During initial configuration steps, MicroExonator extracts annotated microexons and splice sites from one or more gene annotation databases (e.g. GENCODE) and optionally complements them with multiple specialized alternative splicing databases such as VastDB (Tapial et al., 2017). After configuration, SnakeMake enables coordination with cluster schedulers used on high-performance computing platforms or direct process management on a single computer. Thus, given a shortlist of configuration files, MicroExonator can be set to fully reproduce any previous analyses through a single command. Moreover, we provide additional SnakeMake workflows to integrate MicroExonator with downstream quantification steps and to optimize analyses of single cell RNA-seq, which are often much noisier than bulk RNA-seq data.

## Results

### Reproducible detection and quantification of microexons using RNA-seq data

MicroExonator is a computational workflow that integrates several existing software packages with custom python and R scripts to perform discovery and quantification of microexons using RNA-seq data. MicroExonator can analyse RNA-seq data stored locally, but it can also fetch any RNA-seq datasets deposited in the NCBI Short Read Archive or other web-based repositories. As microexon annotations remain incomplete and sometimes inconsistent across different transcript annotations, MicroExonator can incorporate prior information from multiple databases such as RefSeq (Pruitt et al., 2014), GENCODE (Harrow et al., 2006), ENSEMBL (Hubbard et al., 2002), UCSC (Hsu et al., 2006) or VastDB (Tapial et al., 2017). To discover putative novel microexons reads are first mapped using BWA-MEM (Li and Durbin, 2009) to a reference library of splice junction sequences. Misaligned reads are then searched for insertions located at exon-exon junctions. Detected insertions are retained if they can be successfully mapped to the corresponding intronic region with flanking canonical U2-type splicing dinucleotides (Sheth et al., 2006) (Fig 1A). To maximise the number of reads that can be assigned to each splice site, annotated and putative novel microexon sequences are integrated as part of the initial splice tags where they were detected. Reads are re-aligned with Bowtie, performing a fast but sensitive mapping of reads which is further processed to quantify PSI microexon values and perform quantitative filters (Fig 1B, Fig S1).

MicroExonator employs several filters (Fig 1B-C) to remove spurious matches to intronic sequences which may arise due to sequencing errors (Wu and Watanabe, 2005). To illustrate these filters we ran the initial mapping steps over the total RNA-seq from mouse (289 RNA-seq samples from 18 different murine tissues and 1,657 single cells from mice visual cortex (Sloan et al., 2016; Tasic et al., 2016; Weyn-Vanhentenryck et al., 2018)) used in this paper. As a first filtering step, only those insertions that can be detected in a minimum number of independent samples (i.e. technical or biological replicates, three samples is set as default) are considered. Additionally, MicroExonator scores the sequence context of the detected canonical splice sites to measure the strength of their upstream and downstream splice junctions as quantified by a splicing strength score (Parada et al., 2014), and a Gaussian mixture model is used to exclude matches that have weak splice site signals (Fig 1D). Finally, MicroExonator integrates the splicing strength, probability of spurious intronic matching, and conservation (optional) in an adaptive filtering function to remove low confidence candidates (Fig S2).

To ensure that analyses are fully reproducible, MicroExonator was implemented using the SnakeMake workflow manager (Köster and Rahmann, 2012). As MicroExonator may require significant computational resources, SnakeMake also facilitates running on high-performance computer clusters by automating the scheduling of interdependent jobs. SnakeMake itself can be installed from BioConda (Grüning et al., 2018b), and it can initiate MicroExonator directly after downloading the code from our GitHub repository (https://github.com/hemberg-lab/MicroExonator). During runtime, MicroExonator creates custom conda virtual environments which contain specific combinations of software packages found in BioConda repositories to ensure that the same versions are consistently used.

### Benchmarking of computational methods for microexon discovery

To compare MicroExonator with other methods we incorporated a set of synthetic microexons into the GENCODE gene annotation. The microexon sizes were drawn from the previously reported distributions (Irimia et al., 2014; Li et al., 2015) with a greater abundance of in-frame microexons. We also modified the genomic sequence by replacing the intronic flanking regions of simulated microexons with sequences extracted from annotated splice sites. To simulate spurious microexons, we randomly incorporated insertions across splice junctions, as these inserted sequences have the potential to map to intronic spaces.

We used Polyester (Frazee et al., 2015) to simulate reads with a standard Illumina sequencing error rate and processed them using either MicroExonator, Hisat2 (Kim et al., 2015), STAR (Dobin et al., 2013), or Olego (Wu et al., 2013). Our results show that the microexon filtering steps allow MicroExonator to distinguish simulated microexons from spurious microexons with a sensitivity >77% for all microexon lengths (Fig 2A-C). Even though all four aligners could detect a significant fraction of the simulated microexons, they are all limited in their ability to discover very short microexons; STAR’s sensitivity drastically declines for microexons <10 nt, while the sensitivity of Hisat2 and Olego drops for microexons <8 nt. Moreover, the direct output of STAR and Hisat2’s do not represent a reliable source of microexons, as they have low specificity. Using the default parameters results in false discovery rates (FDR) of 43.0% and 33.3%, respectively. Olego had the highest specificity (FDR = 13.0%) of the other mappers, while MicroExonator achieves an FDR of 10.4%. Since MicroExonator’s false discovery events are concentrated in the shortest microexons, discarding microexons <3 nt or <4 nt reduces the FDR to 2.1% and 0.81%, respectively.

**Figure 2.**
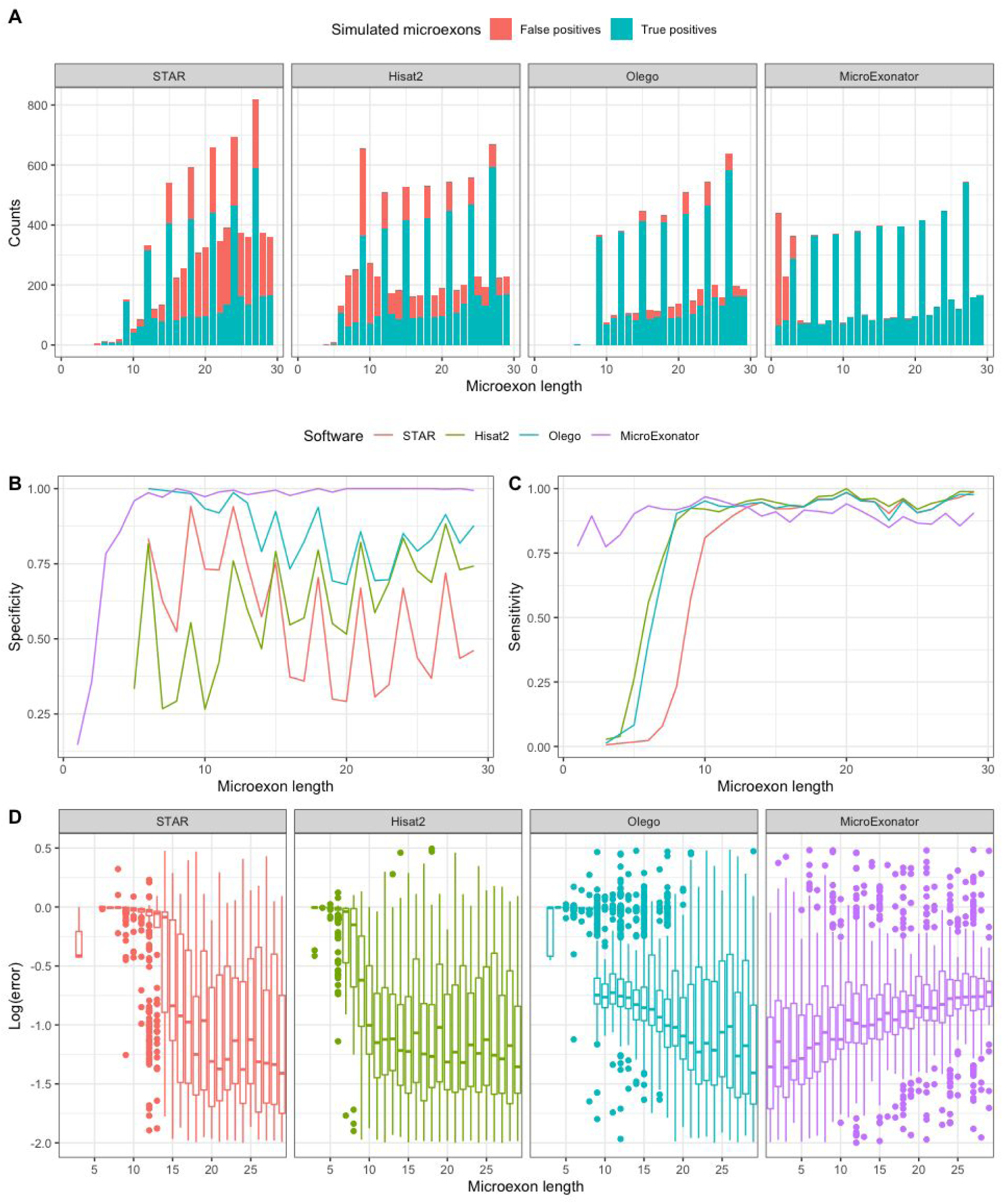
Evaluation of microexon discovery performance of RNA-seq aligners and MicroExonator using synthetic data. A) Size distribution of simulated microexons that were detected by the different software. B-C) Specificity and sensitivity of detected simulated microexons using multiple available tools for evaluation. D) Log10 error PSI values show the accuracy of the microexon quantification.

The simulations also allow us to calculate the ground truth percent spliced in (PSI) values for the microexons, a quantity that represents how frequently a splice junction is incorporated in a transcript. MicroExonator is the only method that has low PSI errors for microexons <10 nt (Fig 2D). Even though MicroExonator’s error rates are slightly higher for microexons >10 nt, they are still comparable to other methods. Taken together, these results show that MicroExonator is more accurate for annotating and quantifying microexons from RNA-seq data compared to conventional RNA-seq aligners.

### Microexon inclusion changes dramatically throughout mouse embryonic development

To investigate how microexon inclusion patterns change during mouse development, we analysed 271 RNA-seq datasets generated by the ENCODE consortium (ENCODE Project Consortium, 2004). These RNA-seq data originate from 17 different tissues, (including forebrain, hindbrain, midbrain, neural tube, adrenal gland, heart, and skeletal muscle) across 7 different embryonic stages (ranging from E10.5 to E16.5), early postnatal (P0) and early adulthood (8 weeks). In addition, we analysed 18 RNA-seq experiments from mouse cortex across nine different time points; embryonic development (E.14.5 and E16.5), early postnatal (P4, P7, P17, P30), and older (4 months and 21 months) (Weyn-Vanhentenryck et al., 2018). Using the annotations provided by GENCODE and VastDB we detected 2,984 microexons in total, and we quantified their inclusion by calculating PSI values for each mouse sample. As some microexons were detected in lowly expressed genes, we only retained microexons whose inclusion or exclusion was supported by >5 reads in >10% of the samples, and this resulted in 2,599 microexons. To characterize the splicing patterns we performed dimensionality reduction using probabilistic principal component analysis (Roweis, 1998; Tipping and Bishop, 1999), and we identified three components that together explain 78.9% of the total PSI variance across samples (Fig 3A-B, Fig S3). The first principal component (PC1) accounts for 56.5% of PSI variance and strongly correlates with the embryonic developmental stage of neuronal samples measured as days post conception (DPC) between E10.5 and E14.5, suggesting strong coordination of microexon splicing during brain embryonic development (Fig 3C, Fig S4). PC2 explains 16.2% of PSI variability and is mostly related to muscular-specific microexon inclusion patterns that were detected in heart and skeletal muscle, suggesting muscle-specific microexon splicing patterns (Fig 3A, Fig S3). Finally, PC3 explains 6.1% of PSI variability and it is related to microexon alternative splicing changes in whole cortex postnatal samples, suggesting that microexon neuronal splicing keeps changing after birth, but to a lesser extent than during embryonic development (Fig 3B, Fig S3).

**Figure 3.**
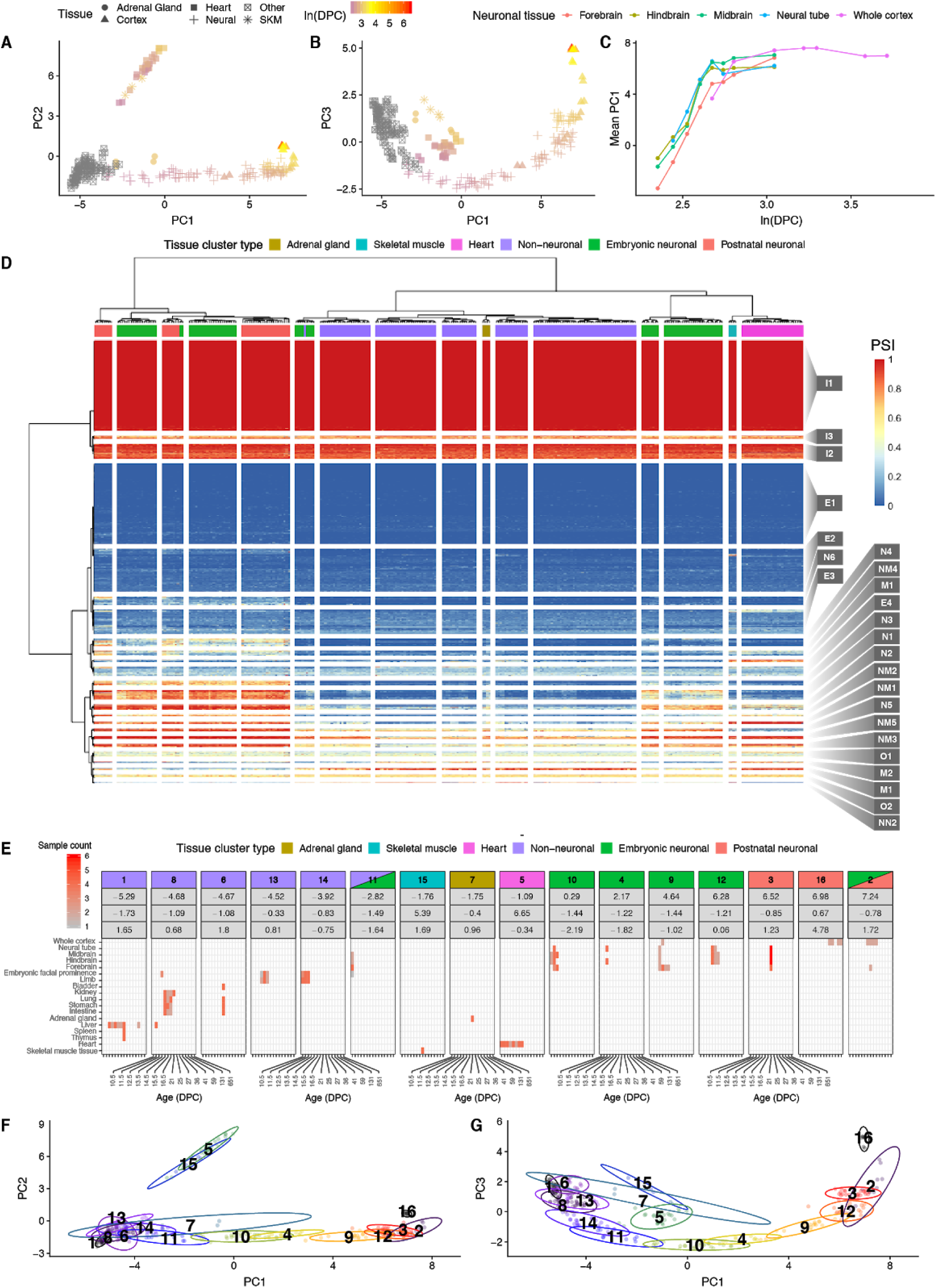
Microexon inclusion through mouse embryonic development. A-B) Dimensionality reduction using probabilistic principal component analysis (PPCA) of microexon PSI values across mouse embryonic and postnatal samples reveals correlation with developmental time for PC1, while PC3 is correlated with developmental time of the postnatal brain samples. Dot shapes denote different sample types and their color indicate their developmental stage, here expressed as log days post conception (DPC). C) PC1 correspondence with embryonic developmental time. D) Heatmap showing microexon inclusion patterns across analysed RNA-seq samples, where rows correspond to microexons and columns to RNA-seq samples. Blue to red colour scale represent PSI values. RNA-seq samples were clustered in 16 different clusters and it composition is colour coded. Microexons were clustered in 24 different clusters and these were named according to their further classification as neuronal (N), muscular (M), neuromuscular (NM), weak-neuronal (WN) and non-neuronal (NN) E) Tissue type and developmental stage composition from sample clusters containing neuronal samples or samples associated with high microexon inclusion. F-G) Projection of the sample clusters across the estimated PPCA components.

To further investigate tissue-specific microexon changes throughout development we performed biclustering of microexon PSI values from the different embryonic samples, and we obtained 24 microexon and 16 sample clusters (Fig 3D). Each of the sample clusters represents a combination of well-defined subsets of tissues and embryonic states (Fig 3E). For example, samples corresponding to brain, heart, skeletal muscles (SKM) and adrenal gland (AG) form separate groups, with the only exception being E10.5 brain samples which clustered together with embryonic facial prominence limb from E10.5 to E12.0. Consistent with the dimensionality reduction analysis, samples from the brain cluster preferentially by developmental time rather than by neuronal tissue, suggesting that microexon alternative splicing changes are greater between developmental stages than between brain regions. As PC1 corresponds to changes of microexon inclusion during neuronal development and PC2 to muscle tissues, we used the mean loading factor values of each microexon cluster from PC1 and PC2 to classify 17 microexon clusters as neuronal (N), muscular (M), neuromuscular (NM), weak-neuronal (WN) and non-neuronal (NN) (Fig 4A-B). Additionally, we found 10 microexon clusters that did not have strong tissue-specific patterns, but were instead either constitutively included (I) or excluded (E) (Fig 4C).

**Figure 4.**
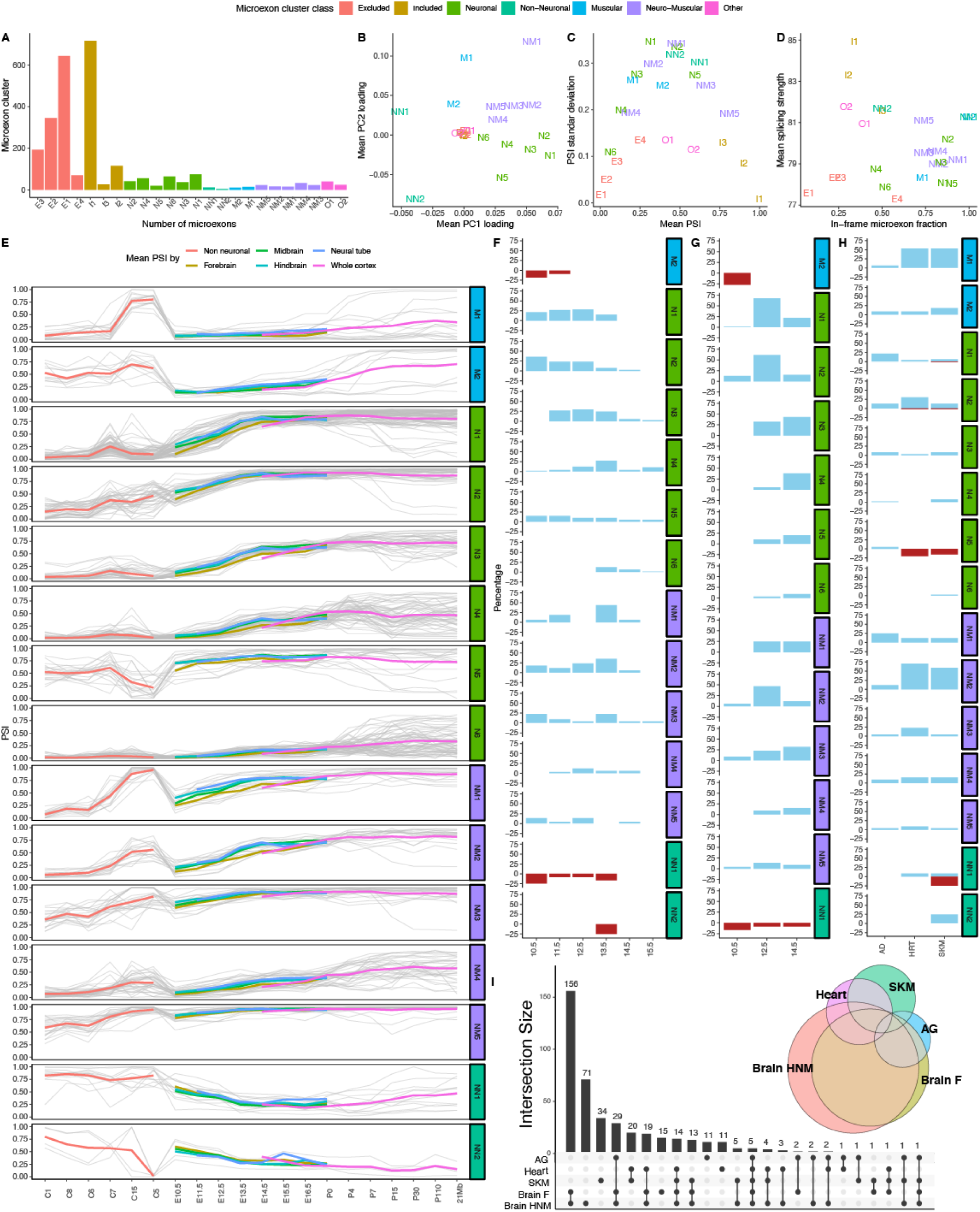
Inclusion properties of microexon clusters. A) Number of microexons belonging to each cluster. B) Mean loading factors for across each cluster for PC1 and PC2. C) Mean and standard deviation of PSI values across microexon clusters. D) Mean U2 scores and in-frame fraction across microexon clusters. E) Mean PSI values across neuronal and neuromuscular microexons. Each grey line represents mean PSI values for a microexon across all samples from a tissue cluster or neuronal developmental stage (x-axis). Colour lines represent mean PSI values across all clusters and stages. F-G) Alternative microexon detected between non-neuronal tissue samples and midbrain, hindbrain and neural tube (F); forebrain (G); adrenal gland (AG), heart (HRT) and skeletal muscle (SKM) (H). Microexon splicing changes are represented percentage of microexons corresponding to each microexon cluster, where microexon inclusion fractions are represented with blue bars and exclusion events with upside down red bars. I) Intersection between microexon sets that were differentially included across sample groups. The vertical bars show the number of microexons corresponding to combinations indicated by the connected dots below. J) Area-proportional Euler diagram representing the most abundant intersections between differentially included microexon sets.

Studies of standard alternative exons have shown that they typically have weaker splice signals than constitutive ones and that they are less likely to disrupt the reading frame (Keren et al., 2010). Thus, we measured the splice site strengths as defined by the average U2 score of microexon flanking splice sites and the fraction of microexons that preserve the reading frame for each cluster (Fig 4D). As expected, the included clusters exhibit the strongest splicing signals, while the excluded clusters have the weakest splice sites, suggesting that constitutive inclusion of microexons relies on strong splicing signals. Moreover, the excluded clusters have a lower fraction of in-frame events, implying that they are likely to be more disruptive to gene function. Interestingly, neuronal, muscular and some neuromuscular clusters have almost as weak splice sites as the excluded clusters, but the total in-frame fraction of these clusters is 0.74. This is considerably higher than the in-frame fractions for longer cassette exons (overall 0.43 and developmentally regulated 0.68) (Weyn-Vanhentenryck et al., 2018). On the other hand, non-neuronal clusters have high U2 scores and also the highest in-frame microexon fraction. The in-frame fraction of each microexon cluster is strongly correlated with the conservation of the coding sequence (Pearson correlation = 0.88, p-value < 1e-7, Fig S5, which implies that microexon clusters with higher conservation tend to preserve the protein frame.

We found a pattern of gradually increased microexon inclusion in the neuronal and neuromuscular categories during mouse brain development in neuronal tissues (Fig 4E, Fig S6). By contrast, non-neuronal microexons exhibited the opposite trend. In addition, neuronal and neuromuscular microexons have higher loading factors on PC1, therefore are the ones might have most variation across mouse embryonic development (Fig S7). To quantitatively assess alternative splicing across mouse brain development, we integrated Whippet (Sterne-Weiler et al., 2018) as part of an optional downstream MicroExonator module. We analysed 221 ENCODE RNA-seq experiments, using 85 non-neuronal samples from the three clusters (C1, C6 and C8) with the lowest PC1 loadings as negative controls. We systematically compared alternative splicing patterns detected in brain, SKM, heart and AG against other non-neuronal tissues. To find microexon splicing changes associated with specific neuronal developmental stages, we pooled by embryonic stage RNA-seq samples from midbrain, hindbrain and neural tube (MHN) between E10.5 and E16.5 and we used Whippet-delta to assess alternative splicing changes using MicroExonator and Whippet PSI values. We found 426 microexons that were consistently detected as differentially included (delta PSI > 0.1 and probability > 0.9 using MicroExonator and Whippet PSI values) in at least one of the comparisons between MHN and controls groups (Fig S8). Interestingly, 326 of these microexon changes are maintained for all subsequent stages once they have been observed. The distribution of the developmental stages when these sustained microexon changes started to be detected differed. While some microexon clusters showed early changes (N1 and N2), other clusters started to be differentially included later on (N3, NM1 and NM2) (Fig 4F). As forebrain tends to show delayed microexon inclusion compared to midbrain, hindbrain and neural tube (Fig 3C, 4E), we pooled forebrain samples between E10.5 and postnatal (P0) and compared samples grouped by developmental stage with the non-neuronal control sample group. We found 407 microexons that were differentially included during at least one forebrain developmental stage, with 257 sustained through all later developmental stages (Fig 4G). In total we found 332 differentially included microexon events that were sustained through all later developmental stages of MHN or forebrain. While all the observed microexon changes across neuronal and neuromuscular clusters correspond to inclusion events, microexons from the non-neuronal cluster (NN1) only correspond to exclusion (Fig 4F-G).

In agreement with previous studies (Irimia et al., 2014; Li et al., 2015) we also found strong inclusion patterns associated with heart and SKM. In addition, we found microexon inclusion patterns associated with AG samples (Fig 3A-B, D-E). Compared with the set of non-neuronal control samples, we found 83, 106 and 58 microexons to be differentially included in heart, SKM and AG respectively (Fig 4E). Most neuronal and neuromuscular microexon clusters show distinct microexon inclusion patterns compared to controls, whereas non-neuronal clusters were associated with microexon inclusion events in heart or exclusion events in SKM samples (Fig 4H).

The set of microexons that were differentially included across the different tissue groups (brain-MHN, forebrain, heart, SKM and AG) overlap. Closer inspection reveals high concordance between the set of microexons associated with sustained changes in inclusion across MHN and forebrain samples. Surprisingly, we found a significant overlap of alternatively included microexons that have concordant patterns in AG and neuronal samples (hypergeometric test p-value < 1e-30). Nearly all of the AG microexons are also found in neuronal samples, but in AG we observed lower PSI values (Fig 4I-J, Fig S9). We hypothesize that the mixture between neuronal and non-neuronal isoforms found in AG is due to the chromaffin cells in the adrenal medulla which are derived from the neural crest and share fundamental properties with neurons (Bornstein et al., 2012; Shtukmaster et al., 2013).

### Microexon alternative splicing is coordinated throughout embryonic development

Based on *in vitro* studies of neuronal differentiation, it has been proposed that microexons are an integral part of a highly conserved alternative splicing network (Irimia et al., 2014). Our analysis of mouse embryonic data (Fig 4E) shows that most microexons remain included once their splicing status has changed. To explore possible functional consequences of these splicing changes we analyzed the interactions between the proteins which contain microexons by constructing tissue specific protein-protein interaction (PPI) networks for brain, heart, SKM and AG using STRING (Szklarczyk et al., 2017). For all four PPI networks the degree of connectivity was significantly higher than expected by chance (p-value<1e-16) (Fig 5A, Fig S10-S11). On average, there were 2.3-fold more connections than expected by chance, with the brain having the largest number of connections (Table S1). Next, we considered the gene ontology (GO) terms and pathways associated with the PPI networks (Fabregat et al., 2018). The Reactome pathways that showed a significant enrichment, include parts of molecular complexes that are involved in membrane trafficking pathways, e.g. “ER to Golgi anterograde transport”, “Clathrin-mediated endocytosis”, “Golgi associated vesicle biogenesis”, “Intra-Golgi and retrograde Golgi-to-ER traffic” and “Lysosome vesicle biogenesis” (Fig 5A-C). We also found a distinct cluster that is annotated as part of “Protein-protein interactions at synapses” (Fig 5D). This group includes presynaptic proteins, e.g. liprins (Ppfia1, Ppfia2 and Ppfia4), protein tyrosine phosphatase receptors (Ptprf, Ptprd, Ptprs) and neurexins (Nrxn1, Nrxn3), which are involved in trans-synaptic interactions with multiple postsynaptic proteins, having a key role in synaptic adhesion and synapse organization. The interactions of these proteins have been shown to be highly regulated by alternative splicing (Takahashi and Craig, 2013), and our results reveal that many of these events occur towards the end of embryonic development (Fig 5F).

**Figure 5:**
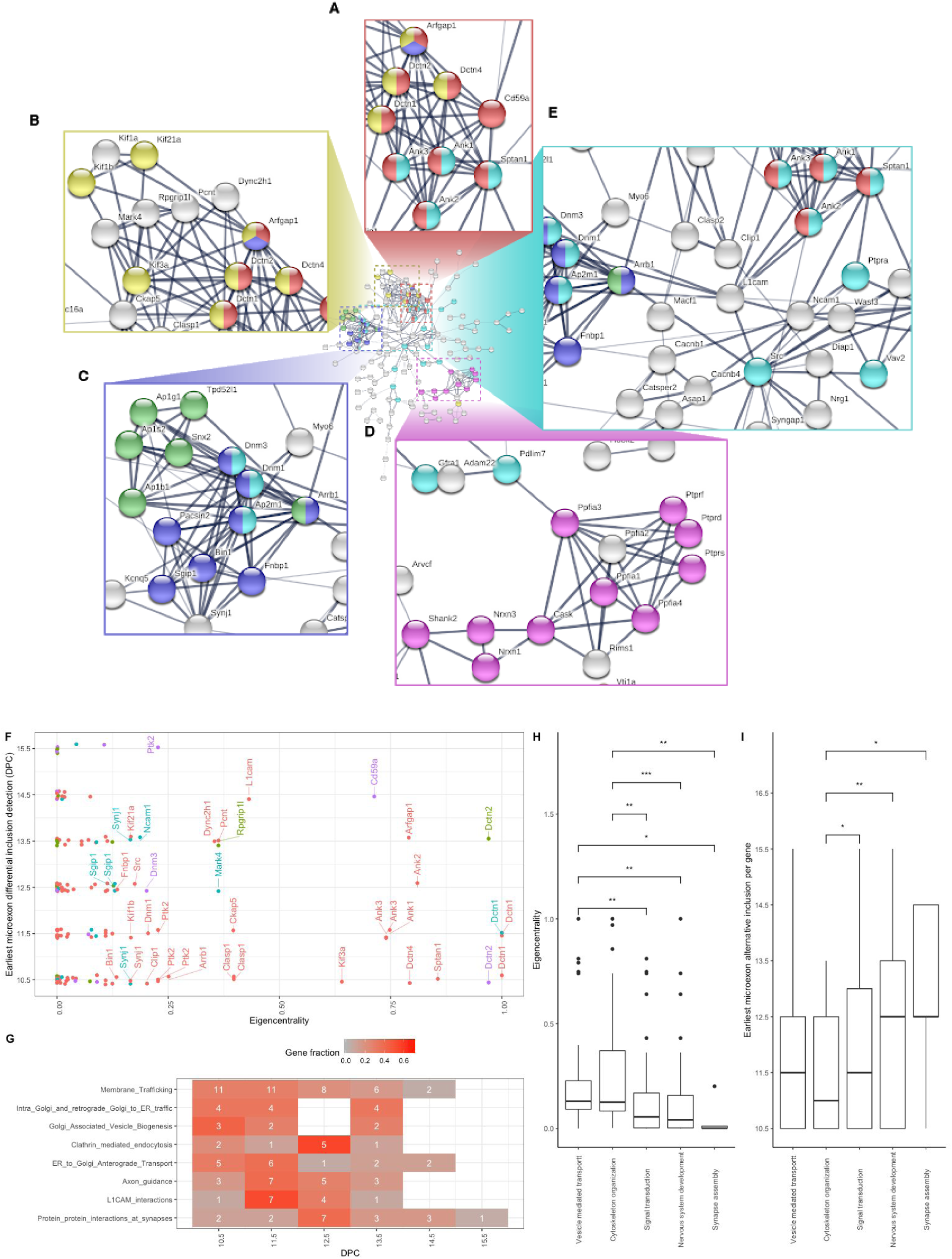
Microexon protein-protein interaction network. A-E) PPI network using as input genes that have microexons that are differentially included across mouse embryonic brain development. Colours represent different Reactome pathways that were enriched on the network; Axon guidance (light blue), Protein-protein interactions at synapses (pink), ER to Golgi anterograde transport (red), Clathrin-mediated endocytosis (dark blue), Golgi associated vesicle biogenesis (green), Intra-Golgi and retrograde Golgi-to-ER traffic (yellow). F) Eigencentrality calculated for each gene node in relation to the developmental stage at which each microexon was included. G) Effect of microexon alternative splicing over different Reactome pathways. Counts indicate the number of microexons that start to be differentially included at each developmental stage for different Reactome pathways that were significant after taking the whole genome as background. H-I) Eigencentrality and earliest developmental stage at which each gene is affected by differential microexon inclusion show differences across some of the GO categories that were significantly enriched after gene background correction. Statistical differences were assessed by one-sided Wilcoxon test while correcting for multiple comparisons. Significant p-values are denoted by * (>0.05), ** (>0.01) and *** (>0.001).

In agreement with previous reports that have highlighted the importance of microexons for axonal and neurite outgrowth (Ohnishi et al., 2017; Quesnel-Vallières et al., 2015), we detected 17 alternative neuronal microexons that affects 14 proteins in the PPI network that are annotated as part of the “Axon guidance” Reactome pathway. These proteins are found in the center of the network and they are connected with the domains involved with membrane trafficking and transsynaptic protein-protein interactions (Fig 5A-E). For two of the proteins associated with this pathway, the non-receptor tyrosine kinase protein Src and L1 cell adhesion molecule (L1cam), microexon inclusion is known to play a key role in neuritogenesis (Kamiguchi and Lemmon, 1998; Keenan et al., 2017), but the importance of microexons in other proteins in this pathway remains poorly characterised. At early developmental stages (E10.5-E11.5) we found several microexon alternative splicing events in genes associated with “membrane trafficking” pathways concentrated. A subset, “clathrin mediated endocytosis” is associated with microexon changes in the later stages, as most events became significant only after E12.5 (Fig 5G). Similarly, “axon guidance” microexon changes mostly occur at E11.5, in particular the microexon alternative splicing events for proteins that interact with L1cam.

Since microexon inclusion occurs in several waves (Fig 4E), we hypothesized that the temporal dynamics would be reflected in the topology of the PPI network. To quantitatively evaluate the position of each gene in the network, we calculated several centrality measures. The result is not straightforward to interpret since several of the central nodes feature more than one microexon inclusion event (e.g. Synj1, Ank3 and Dctn2), which sometimes emerge at different embryonic stages (Fig 5F). Nevertheless, the results show that L1cam and 7 out of 10 of its interactors are amongst the 25% of nodes with highest eigencentrality and that Src has the highest harmonic centrality and betweenness. An investigation of genes corresponding to some of the most relevant GO terms revealed that the genes with the highest eigencentrality are the first to be included (Kruskal-Wallis rank sum, p-values < 0.05) (Fig 5H-I).

### MicroExonator enables the identification of novel neuronal microexons

Of the 332 microexons that were differentially included across brain development, 98 were inconsistently annotated as compared to GENCODE and VastDB. Of these 98 neuronal microexons, we found 35 that are only annotated in GENCODE, and 30 neuronal microexons that are not annotated in GENCODE, but are present in VastDB. Despite the fact that the mouse genome is comprehensively annotated, we found 33 neuronal microexons that are not annotated in GENCODE nor VastDB. Due to the high sensitivity and specificity demonstrated in simulations (Fig 2), we expect that all 31 microexons ≥6 nts are true positives (Fig 6A).

**Figure 6.**
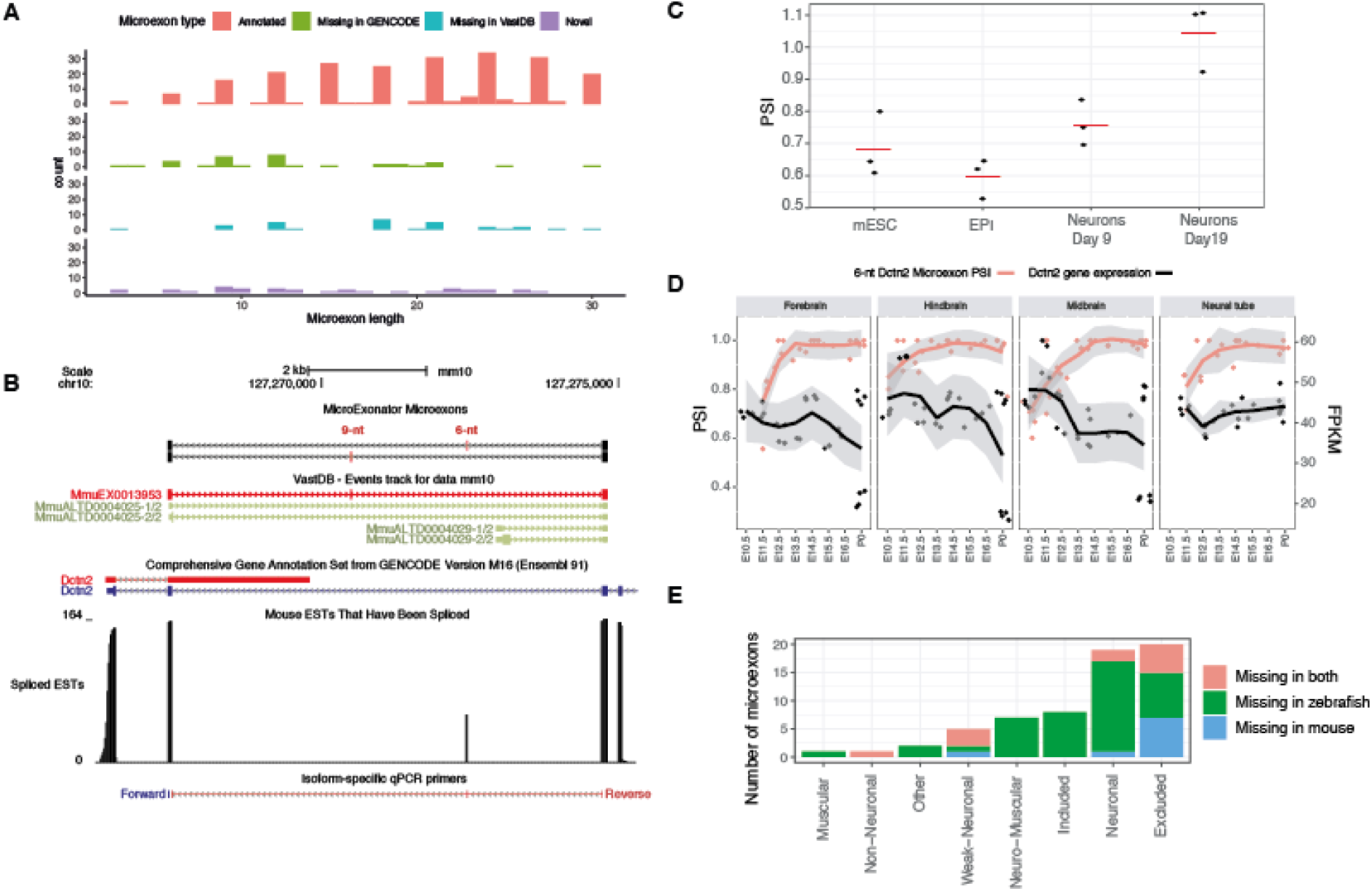
Discovery of novel microexons in mouse and zebrafish. A) Histogram showing size distribution of microexons that were found to be differentially included across mouse embryonic development. B) Alternative Dctn2 microexons that are inconsistently annotated in mouse GENCODE and VastDB annotations. C) Novel 6-nt Dctn2 microexon shows a progresive inclusion through mouse embryonic development. D) PSI values calculated from normalized RT-PCR measurements shows a gradual inclusion of the 6-nt Dctn2 microexon though *in vitro* neuronal mESC to neuron differentiation. E) Number of conserved microexons between mouse and zebrafish that are missing from their transcript annotation.

To validate one of the novel microexons, we focused on the Dctn2 gene (eigencentrality of 0.76), where we detected two adjacent differentially included microexons of length 9 and 6 nts (Fig 6B). Neither of these microexons are annotated in GENCODE, but the 9-nt microexon is annotated in VastDB (MmuEX0013953). Interestingly, the downstream 6-nt microexon that was discovered by MicroExonator is validated by spliced ESTs (Benson et al., 2004). We detected differential inclusion of the 6-nt Dctn2 microexon from E10.5 in MHN samples, whereas in forebrain it is differentially included from E12.5 (Fig 6C).

We performed qRT-PCR experiments to assess the inclusion of the Dctn2 6-nt microexon during a mESC to neuron differentiation protocol using one set of primers that were designed to amplify Dctn2 isoforms with 6-nt microexon inclusion and another set to amplify total Dctn2 isoforms. Next, we calculated the ratio of 6-nt inclusion across mESC, EPI cells and differentiated neurons at two different stages (Fig 6D). The inclusion ratios from the qRT-PCR measurements indicate that the Dctn2 6-nt microexon is included through *in vitro* differentiation of mESC to neuron, consistent with our findings during embryonic development for this microexon. These results show that the alternative splicing quantification provided by MicroExonator can identify novel microexons, even for model organisms that are well annotated.

### Identification of microexons in zebrafish brain

To demonstrate how MicroExonator can be applied to species with less complete annotation, we analyzed 23 RNA-seq samples from zebrafish brain (Park and Belden, 2018). We found 1,882 microexons, of which 23.8% are not found in the ENSEMBL gene annotation. We used liftover (Hinrichs et al., 2006) to assay whether some of these microexons are evolutionary conserved microexons in mouse, and we successfully mapped 401 zebrafish microexons. Of these, 85% mapped directly to a previously identified mouse microexon, and most of the remaining 15% mapped to longer exons. Mapping the microexons in the other direction, 617 out of the 2,938 that were identified from the mouse development data mapped to the zebrafish genome and 49.7% of those in return mapped to a zebrafish microexon. By integrating these results we obtained a total of 402 microexons pairs that are found in both zebrafish and mouse. Since 90.3% of the pairs had identical length in both species, they are highly likely to correspond to evolutionary conserved microexons.

To compare the microexon annotation between mouse and zebrafish, we asked how many of the 402 conserved microexons that are missing in mouse or zebrafish gene transcript annotation. While only 6.9% of these exons are missing from the mouse transcript annotation provided by GENCODE, 16.1% are missing from the ENSEMBL zebrafish transcript annotation. Moreover, the largest fraction of conserved microexons that are missing in zebrafish transcript annotation corresponds to neuronal microexons (Fig 6D).

### Cell type specific microexon inclusion in mouse visual cortex

Our analysis of neuronal development suggested that the main difference in microexon inclusion is between time points rather than tissues. However, since these data do not reflect the diversity of cell types within neuronal tissues, we hypothesized that microexon inclusion patterns may vary amongst different neuronal subtypes. To study cell-type specific patterns of microexon inclusion, we analyzed SMART-seq2 scRNA-seq data from visual cortex of adult male mice (Tasic et al., 2016). The sample contains 1,657 cells which were assigned into six cell type classes that were further subdivided into 49 distinct clusters.

We focused on the GABA-ergic and the glutamatergic clusters of neurons which contain 739 and 764 cells respectively. We first ran the microexon discovery module with an expanded annotation, which included the microexons discovered from our previous analyses. This yielded 2,344 microexons that were included in at least one cell. Next, we used Whippet to quantify the PSI of the microexons detected by MicroExonator for each cell. Due to the sparse coverage, the single-cell analysis is sensitive to errors. Thus, for each neuronal type we also pooled 15 randomly selected neurons into pseudo-bulk groups that were quantified by Whippet. When we processed PSI measurements at the single cell level, we obtained 19 microexons that were differentially included between GABA-ergic and glutamatergic neurons. The analysis of pseudo bulk PSI values identified a total of 38 differentially included microexons, 20 of which were also identified from the single cell PSI values (Fig S12, Table S2).

Among the genes that contain differentially included microexons between GABA-ergic and glutamatergic neurons is a group of eleven genes that encode for proteins that localize at synaptic compartments. We found seven presynaptic proteins, two postsynaptic proteins and two proteins that have been observed at both locations (Fig 7A). For example, the type IIa RPTPs subfamily of proteins undergoes tissue-specific alternative splicing that determines the inclusion of four short-peptide inserts, known as mini-exon peptides (meA-meD) (Pulido et al., 1995a, 1995b; Takahashi and Craig, 2013). While meB comprises four residues (ELRE) and is encoded by a single microexon, meA has three possible variants that can form as a result of the combinatorial inclusion of two microexons; meA3 (ESI), meA6 (GGTPIR) and meA9 (ESIGGTPIR) (Yamagata et al., 2015a). Our analysis shows a consistent inclusion of meB in both GABA-ergic and glutamatergic neurons. However, we detected cell type specific rearrangement of meA microexons which promotes inclusion of meA9 in glutamatergic neurons, while in GABA-ergic neurons meA variants are mostly excluded (Fig 7B). Alternative splicing of meA/B microexons are key to determining the selective trans-synaptic binding of PTPδ to postsynaptic proteins, which is a major determinant of synaptic organization (Takahashi and Craig, 2013). In addition, we found other alternatively spliced microexons in genes that are involved in synaptic cell-adhesion, e.g. Gabrg2, Nrxn1 and Nrxn3 (Südhof, 2017; Takahashi and Craig, 2013). The microexon inclusion in these genes is variable across the core clusters, sometimes showing stark differences between GABA-ergic and glutamatergic neuron subtypes (Fig 7C). These results suggest that microexon inclusion is not only coordinated at the tissue-type level, but that it is also finely tuned across neuronal cell-types, and these differences may be of importance for determining neuronal identity and synapse assembly.

**Figure 7.**
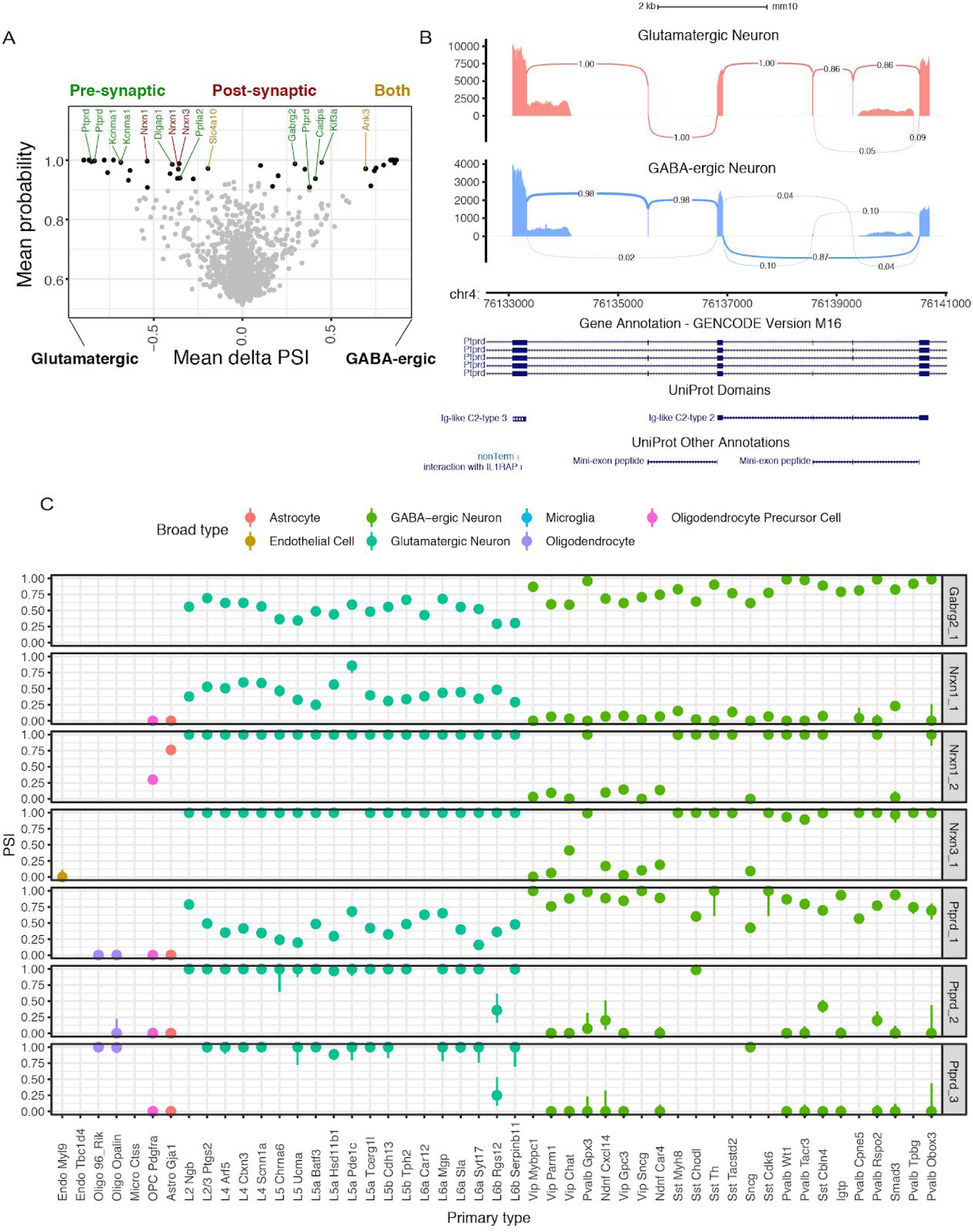
Differential alternative splicing analysis of microexons between glutamatergic and GABA-ergic neurons using scRNA-seq data. A) Volcano plot showing alternatively included microexons between glutamatergic and GABA-ergic neurons. Differentially included microexons are highlighted in black. Synaptic proteins are labeled with different colours depending on their sub-synaptic localization. B) Sashimi plot showing PTPδ microexons that determine the inclusion of meA/B mini-exon peptides. C) Microexon inclusion patterns across all core clusters at proteins involved in trans-synaptic interactions.

## Discussion

We have presented MicroExonator, a complete bioinformatic workflow for reproducible discovery and quantification of microexons. MicroExonator is designed to handle large volumes of data and it will automatically download datasets and schedule jobs on a computer cluster. Currently, it is the only publicly available method that allows these types of investigations. MicroExonator’s discovery module is based on detection of inserted sequences between annotated splice sites which enables the identification of very short microexons that cannot be reliably detected by spliced RNA-seq aligners (Fig 2C). Thus, MicroExonator will greatly facilitate the study of microexons. MicroExonator is straightforward to incorporate into an existing RNA-seq analysis workflow. Importantly, MicroExonator can be used to directly study microexon conservation across species, thereby making it possible to understand if inclusion patterns are as well conserved as the nucleotide sequences. Furthermore, MicroExonator makes it possible to study RNA-seq data from large cohorts to investigate if there are microexons that differ amongst individuals.

As proof of principle, we used MicroExonator to analyse RNA-seq data from 301 RNA-samples from mice at embryonic and adult stages and 1,679 single cells. We have expanded the catalog of murine microexons by identifying 941 previously uncharacterized loci. In agreement with previous analyses, we identified microexons that were differentially included in brain, heart and SKM (Irimia et al., 2014; Li et al., 2015), but we also detected 58 microexons that are differentially included in adrenal gland. Taken together, we have presented the most comprehensive catalog of microexons available to date, and it allowed us to uncover several distinct inclusion patterns in both developing and adult mice.

### Microexon coordination across neuronal development

Our quantitative analysis revealed that the proteins containing microexons form a highly connected network during mouse neuronal development. Moreover, analysis of the topology of the network suggests that the microexons for the most central nodes are included early in development. It is not yet fully understood how this coordination is achieved, but it has been proposed that microexon inclusion relies on an upstream intronic splicing enhancer which is recognized by specific neuronal splicing factors (Gonatopoulos-Pournatzis et al., 2018). However, we also identified a large group of microexons that are constitutively included across murine tissues, suggesting that their inclusion cannot be dependent on tissue-specific factors alone. Instead, our analysis points to a more straightforward explanation as the constitutive microexons have stronger splicing signals than neuronal microexons. Further analysis of neuronal microexon cis-regulatory elements is required to understand how inclusion events are coordinated and why there there is a small number of microexon that is progressively excluded through brain development.

The predominant mechanism for regulating alternative splicing events during neuronal development is through RNA binding proteins (Vuong et al., 2016). In the case of microexons, SRRM4 and RBFOX1 have a critical role in coordinating microexon inclusion through brain development, and changes in expression of these splicing factors has been linked to misregulation of alternative splicing events in individuals with autism spectrum disorder (ASD) (Irimia et al., 2014; Li et al., 2015; Voineagu et al., 2011). In fact, alternative splicing changes associated with ASD are enriched in microexons and they are recapitulated in mutant mice haploinsufficient for SRRM4 (Irimia et al., 2014; Quesnel-Vallières et al., 2015). Moreover, a recent genome-wide CRISPR-Cas9 screen has identified two additional factors, SRSF11 and RNPS1, that contribute to SRRM4-dependent microexon regulation, and these genes have also been implicated in ASD and other neurological disorders (Gonatopoulos-Pournatzis et al., 2018). Another example of a protein where imbalances of microexon inclusion has been associated with elevated risk of ASD is cytoplasmic polyadenylation element binding protein 4 (CPEB4) (Parras et al., 2018). We found differential inclusion of CPEB4 microexon during mouse embryonic brain development, and we also found microexon changes in other protein factors that are involved in mRNA polyadenylation, such as CPEB2, CPEB3 and FIP1L1. However the role of these microexons in neuronal function and neuropsychiatric diseases remains unexplored.

The high degree of conservation of microexons strongly suggests that they are functionally important, but for the most part we lack a detailed, mechanistic understanding. A notable exception is Src where microexon inclusion leads to the production of n-Src, a well characterized neuronal splice variant. The Src microexon encodes for a positively charged residue located at an SH3 domain that has been shown to regulate Src kinase activity and specificity (Brugge et al., 1985). From the STRING analysis we found evidence for Src-dependent phosphorylation of Git1, Ctnnd1 and Ptk2 (Chernyavsky et al., 2008; Lim et al., 2002; Wang et al., 2010), though the impact of neuronal microexon alternative Forsplicing for these phosphorylation events remains unknown. Moreover, recent studies show that n-Src microexon inclusion is required for normal primary neurogenesis and L1cam dependent neurite elongation (Keenan et al., 2017; Lewis et al., 2017), implying a strong phenotype. Another central node in the PPI network that is known to undergo microexon alternative splicing changes that are important for axon growth is L1cam, a founding member of L1cam protein family. Across the L1cam protein family a sorting signal is included due to 12-nucleotide alternative microexons. In the case of L1cam, the 12-nucleotide microexon mediates its clathrin mediated endocytosis by interacting with adaptor protein complex 2 (AP-2) (Kamiguchi et al., 1998). Our analysis shows that the AP-2 mu subunit (Ap2m1) is also affected by microexon inclusion through mouse brain development.

### Cell-type specific microexon alternative splicing across the mouse visual cortex

Single cell RNA-seq data is providing an unprecedented opportunity to survey cell-specific expression profiles. However, with a few notable exceptions (Arzalluz-Luque and Conesa, 2018; Gokce et al., 2016; Lukacsovich et al., 2019; Zhang et al., 2016), most scRNA-seq analyses have focused on analysis at gene rather than the transcript level. Here, we applied MicroExonator to GABA-ergic and glutamatergic cells from visual cortex, and to increase the power we developed a downstream SnakeMake workflow, snakepool. As many splicing events are undetected in single cell data due to poor coverage, a pooling strategy is necessary to increase the power to identify significant differential inclusion events.

We identified 39 microexons that were differentially included between GABA-ergic and glutamatergic neurons and 11 synaptic proteins that are affected by 15 of these cell-type specific microexons. Among these proteins, we found two alternatively included microexons on Ptprd, which are known to have a key role in modulating trans synaptic interactions, thus having a direct impact on synapse formation (Yamagata et al., 2015a, 2015b). In addition we also show that microexons found in Ptprd and other proteins involved in transsynaptic protein interactions can have distinctive alternative inclusion profiles across GABA-ergic and glutamatergic subtypes (Fig 7C). Importantly, this result demonstrates that even though bulk RNA samples from different brain regions are largely similar, there are differences between both neuronal and non-neuronal populations. The differential inclusion of microexons could have profound effects on neuronal identity, synapse formation and disease. For example, the differential microexon inclusion event that we identified in GABAa receptor subunit γ (GABRG2) can have a direct impact on GABA-ergic neurons as this microexon introduces a phosphorylation site that regulates GABA activated current. Misregulation of this alternative splicing event has been associated with schizophrenia in human patients (Ustianenko et al., 2017). However, additional analyses alternative microexon patterns across neuronal cell-types will be required to fully understand their contribution to neuronal heterogeneity and function.

## Acknowledgements

We would like to thank Nadav Ahituv, Chris Smith, Christoph Schlaffner and Judith Steen for helpful feedback on the manuscript. Also, we thank David Jordan, Gero Bohner, Lorenzo Loyola, Simon Bajew, Tallulah Andrews and Vladimir Keselev for relevant discussion during the development of this project.

## Author Contributions

GEP, RM, EAM and MH conceived the project. GEP wrote the code. GEP and IGS analysed the data. HF carried out the qRT-PCR validation experiments. EM, MAE, EAM and MH supervised the research. GEP and MH wrote the manuscript with input from all other authors.

## Methods

### Annotation guided microexon discovery using RNA-seq data

MicroExonator was implemented over the Snakemake workflow engine (Köster and Rahmann, 2012), to facilitate reproducible processing of large numbers of RNA-seq samples. During an initial discovery module, MicroExonator uses annotated splice junctions supplied by the user (a gene model annotation file can be provided in GTF or BED format) to find novel microexons. RNA-seq reads are first mapped to a library of reference splice junction tags using BWA-MEM (Li and Durbin, 2009) with a configuration that enhances deletion detection (bwa mem -O 6,2 -L 25). The library of splice junction tags consists of annotated splice junctions between exons ≥ 30 nt and spanning introns ≥80 nt. For each splice junction, a reference sequence tag is generated by taking 100 nt upstream and downstream from the corresponding transcript sequence. Splice junction alignments are processed to extract read insertions with anchors ≥8 nt that map to exon-exon junction coordinates. Inserted sequences are then re-aligned inside the corresponding intronic sequence, but only matches flanked by canonical splice site dinucleotides (GT-AG) are retained (Figure 1A). The obtained reads are re-mapped to the reference genome using hisat2 (Kim et al., 2017). A preliminary list of microexon candidates is generated based on reads whose insertions are aligned to the intronic spaces with no mismatches, and that could not be fully mapped to the reference genome (soft clipping alignments are ignored).

### Quantification of microexon inclusion

In a subsequent quantification module, novel microexon candidates are integrated into the gene annotation to generate a second library of splice junctions tags, where putative novel loci from the discovery phase and annotated microexons are integrated at the middle of the tag sequences (Figure 1B). Reads are aligned again to this expanded library of splice junction tags using Bowtie (Langmead et al., 2009), which performs a fast ungapped alignment allowing for 2 mismatches (bowtie -v 2 -S). Reads that map to the splice junction tags are also mapped to the reference genome using Bowtie, also allowing two mismatches. Reads that could only fully map to a single splice junction tag but no other location count towards novel or annotated microexons.

### Filtering of spurious intronic matches

MicroExonator uses a series of filters to distinguish real splicing events that may result in a novel microexon of length *L* from spurious matches with intronic sequences. Since we only allow for intronic matches that are flanked by canonical dinucleotides (4 nt), we search for matching sequence of length *L + 4* in the intron.

For a random sequence of length *L*, where all four nucleotides have the same frequency, the probability of at least one spurious match inside an intron with flanking GT-AG dinucleotides, *P_s_*, can be calculated as *P_s_ = 1 - (1 - 1/4^L+4^)^K^*, where *K* is the number of k-mers of length *L + 4* that are found in an intron of length *N*, with *K = N - L - 4*. Since microexons shorter than 3nt cannot be identified with high specificity, they are reported as a separate list without further filtering.

Microexons that are 3 nt or longer are filtered further by evaluating the splice site signal by measuring the match to the canonical splicing motif as defined by the U2-type intron position frequency matrices (Sheth et al., 2006). We normalize the score to range from 1 to 100, and we call this quantity U2 Score. We then fit the distribution of U2 scores using a two component Gaussian mixture model (Figure 2B), and from this we calculate a score, *M_s_*, for each putative microexon as *M_s_ = 1 - (1 - P_s_P_U2_)/n*, where *P_U2_* is the probability that the observed U2 score came from the Gaussian with the higher mean and *n* is the number of matches for a given intron. Finally, MicroExonator calculates an adaptive threshold, *M\**, to determine the minimal *M_s_* score required. Let *R* denote the number of detected microexons that have *M_s_>t*. A linear model is used to fit *R*^t^ as a function of their length, with *t* ranging between 0 and 1. MicroExonator recommends *M\** as the score corresponding to *t\**, the value which results in the minimal residual standard deviation sum. This threshold is used to generate a high confidence list of microexons, but all detected microexon are reported across different output files. By default, MicroExonator uses *M\** to filter out microexons with low scores, but the threshold can be set manually by the user. If conservation data (e.g. Phylop/Phastcon) is provided, then microexons with *M_s_<M* that exceed a user-defined conservation threshold (default value = 2) are also included in a high confident list of microexons and flagged as “rescued”.

### RNA-seq simulation

We used Polyester (Frazee et al., 2015) to simulate RNA-seq reads from modified mouse GENCODE gene models (V11). To generate true positive microexons, we inserted a set of 4,930 randomly selected sequences with a length ranging from 1 to 30 nucleotides inside annotated introns longer than 80 nts. At the same time we swapped the original intronic sequences of annotated microexons for splicing signals found at another randomly selected annotated exon. To simulate spurious microexon matches (false positive microexons), we randomly included 5,180 insertions corresponding to intronic sequences at exon-exon junctions that were not flanked by canonical splicing sequences. The insertion rates and lengths were simulated using parameters extracted from real RNA-seq experiments from postnatal forebrain samples (Fig S1C). Our simulations provide a realistic set of false positive microexons that emulates real RNA-seq experiment condition as closely as possible.

### Microexon analyses across mouse development using bulk RNA-seq data

As a proof of principle, we applied MicroExonator to 283 RNA-seq datasets obtained from ENCODE project (Sloan et al., 2016), corresponding to embryonic and postnatal tissue samples coming from 17 different tissues. We used mm10 mouse genome assembly obtained from UCSC genome browser database (Karolchik et al., 2003), and as source of annotated splice junctions we used the union of GENCODE Release M16 (Harrow et al., 2006) and VastDB (Tapial et al., 2017). We quantified novel and annotated microexons through Percent of spliced-in (PSI) values by using MicroExonator’s built-in scripts, or by using Whippet (Sterne-Weiler et al., 2018). Bi-clustering of samples and microexons was performed by applying the Ward’s minimum variance criterion in R (Müllner and Others, 2013; Murtagh and Legendre, 2014) over a MicroExonator Euclidean distance matrix where the similarity of the samples was calculated from the PSI values (Supplemental Methods - R Notebook file).

PSI values were also used to perform PPCA using the ppca function from the pcaMethods R library (Stacklies et al., 2007). The obtained PPCA loading factors were used to classify microexon clusters. Assuming that PC1 and PC2 are related with variance observed at brain and muscle respectively, loading factors can be used to evaluate the tissue-specificity of microxon inclusion. Thus, microexons that have loading factors >0.03 for PC1 and PC2, were considered as neuromuscular (NM1-3). The ones that only have high loading factors for either PC1 or PC2 were considered as neuronal (N1-4) and muscular (M1-3), respectively. We found one microexon cluster with a significant negative loading factor over PC1 (<-0.03), which we considered to be non-neuronal (NN1). We also found microexon clusters that have a consistent inclusion (I1-7) or exclusion (E1-5) pattern across all samples.

We quantified splicing nodes using Whippet’s quantification module (whippet-quant.jl) and we supplied MicroExonator output as input to the Whippet differential inclusion module (whippet-delta.jl). We used both MicroExonator and Whippet quantification to assess changes in microexon inclusion between different sample groups. A baseline was defined by the non-neuronal tissue clusters that had the lowest PC1 values (clusters C1, C6 and C6). Different neuronal sample groups were defined by developmental stages (E10.5, E11.5, E12.5, E13.5, E14.5, E15.5) and brain tissue type; samples corresponding to midbrain, hindbrain and neural tube were pooled together (MHN sample group) whereas forebrain samples were evaluated independently. Moreover, additional non-neuronal groups were formed according to their tissue of origin, which correspond to heart, SKM or adrenal gland. Each sample group was compared with the baseline groups. Across the different comparisons, we only considered as significant those microexons which have >0.9 probability of being differentially included and ≥0.1 delta PSI values. To further avoid quantification errors, we only selected those microexons that were detected as differentially included using both MicroExonator and Whippet quantification. The sets of genes that have at least one microexon differentially included in brain, SKM, heart or adrenal gland were analyzed by building a protein-protein interaction network using STRING (Szklarczyk et al., 2017). PPI network analyses were performed using STRING v.11.0 (Szklarczyk et al., 2017), through the main web server (https://string-db.org/) taking as input the ENSEMBL ID of the set of genes which contains one or more microexons.

### Neuronal mouse dopamine neuron preparation and RT-PCR validations

Mouse embryonic stem cells (mESC) were differentiated into dopamine neurons as previously described (Metzakopian et al., 2015). Briefly, mESCs were first differentiated into Epiblast stem cells (EPI) using fibronectin coated plates and N2B27 basal media (composed of Neurobasal media, DMEM/F12, B27 and N2 supplements, L-glutamine and 2-Mercaptoethanol) supplemented with FGF2 (10mg/ml) and Activin A (25mg/ml). After three passages, EPI were differentiated into dopaminergic neurons using plates collated with poly-L-lysine (0.01%) and Laminin (10ng/ml) and N2B7 media supplemented with PD0325901 (1mM) for 48hours (Day 0 to Day 2). 3 days later (Day 5), N2B27 media was supplement with Shh agonist SAG (100nM) and Fgf8 (100ng/ml) for 4 days. Media was then changed to N2B27 media supplemented with BDNF (10ng/ml), GDNF (10ng/ml) and ascorbic acid (200nM) from Day 9 onwards. During neuronal differentiation cells were passaged at Day 3 and Day 9. Cells were collected for qRT-PCR analysis at several stages: mESC, EPI, Day 9 neurons and Day 19 neurons. RNA extraction was performed using the RNeasy Mini Kit (Qiagen) and samples analysed with a QuantStudio 5 PCR system (Thermo Fisher Scientific).

### Microexon identification in Zebrafish brain

RNA-seq experiments for Zebrafish brain tissues were obtained from (Park et al., 2018) using GEO accession code GSM2971317. Microexon detection and quantification were performed with MicroExonator using default parameters based on Ensembl gene predictions 95 and the danRer11 genome assembly. To compare mouse and zebrafish microexons, we performed a batch coordinate conversion using the liftOver script from UCSC utilities (Karolchik et al., 2003).

### Single cell analyses

We applied MicroExonator to single cell RNA-seq data from mouse visual cortex (Tasic et al., 2016). In addition to MicroExonator PSI quantification, we also computed PSI using Whippet for all detected microexons found. To compare microexon inclusion rates, we used Whippet to perform an iterative quantification and inclusion analysis. We pooled data coming from neuronal cell-types; GABA-ergic and glutamatergic in pseudo-bulk groups of 5 cells (or fewer for the last group), and repeated this process 10 times. During each iteration, splicing node PSI values were calculated using whippet-quant.jl. Both single-cell and pseudo-bulk were used to assess differential inclusion of splicing nodes. Only those microexons with >0.9 mean probability and where the difference in delta PSI between pooled and unpooled analysis was <0.25, were included. Sashimi plots were generated by adapting ggsashimi (Garrido-Martín et al., 2018) to display the total number of reads that is supported by each splice site (Supplemental Material). The read counts were subsequently used to calculate splice site usage rates.

## Software availability

MicroExonator is available at https://github.com/hemberg-lab/MicroExonator (under MIT licence), together with additional post-analysis code.

## Supplemental Material

**Figure S1:**
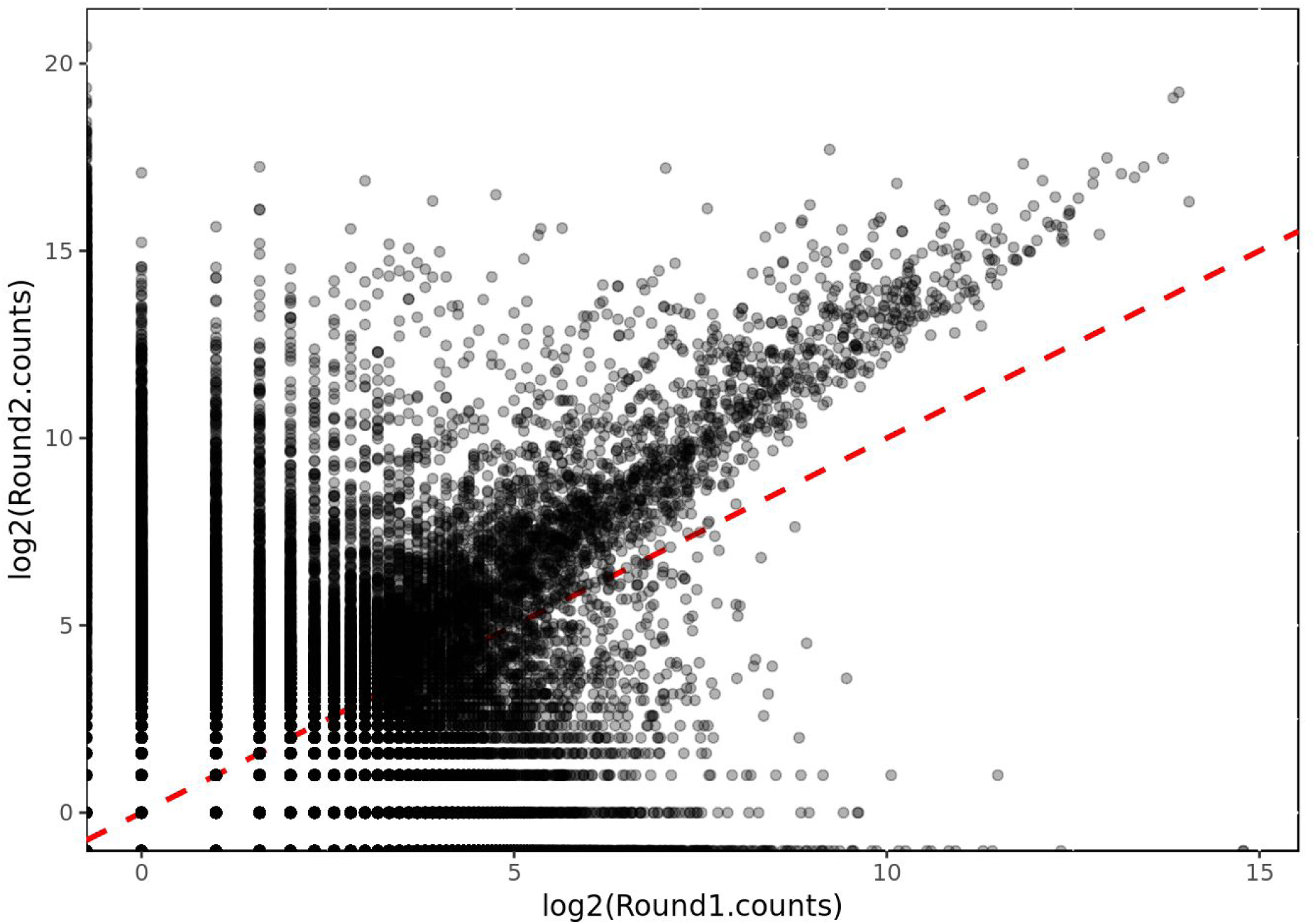
Number of reads assigned to microexon splice sites during the first and second splice junction alignment.

**Figure S2.**
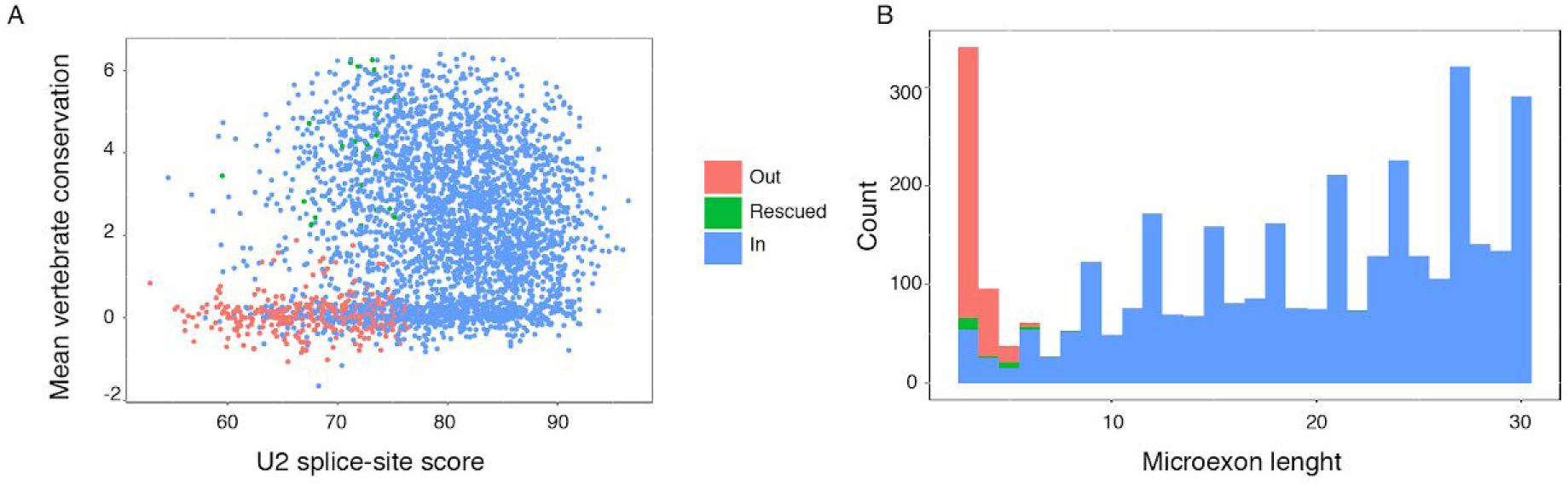
Filtering of putative novel and annotated microexons. A) Conservation and splicing strength of microexons that pass the final quantitative filtering steps. A few microexons were initially filtered out were rescued based on their conservation signal (Phylop score >= 2). B) Size distribution of filtered microexons.

**Figure S3.**
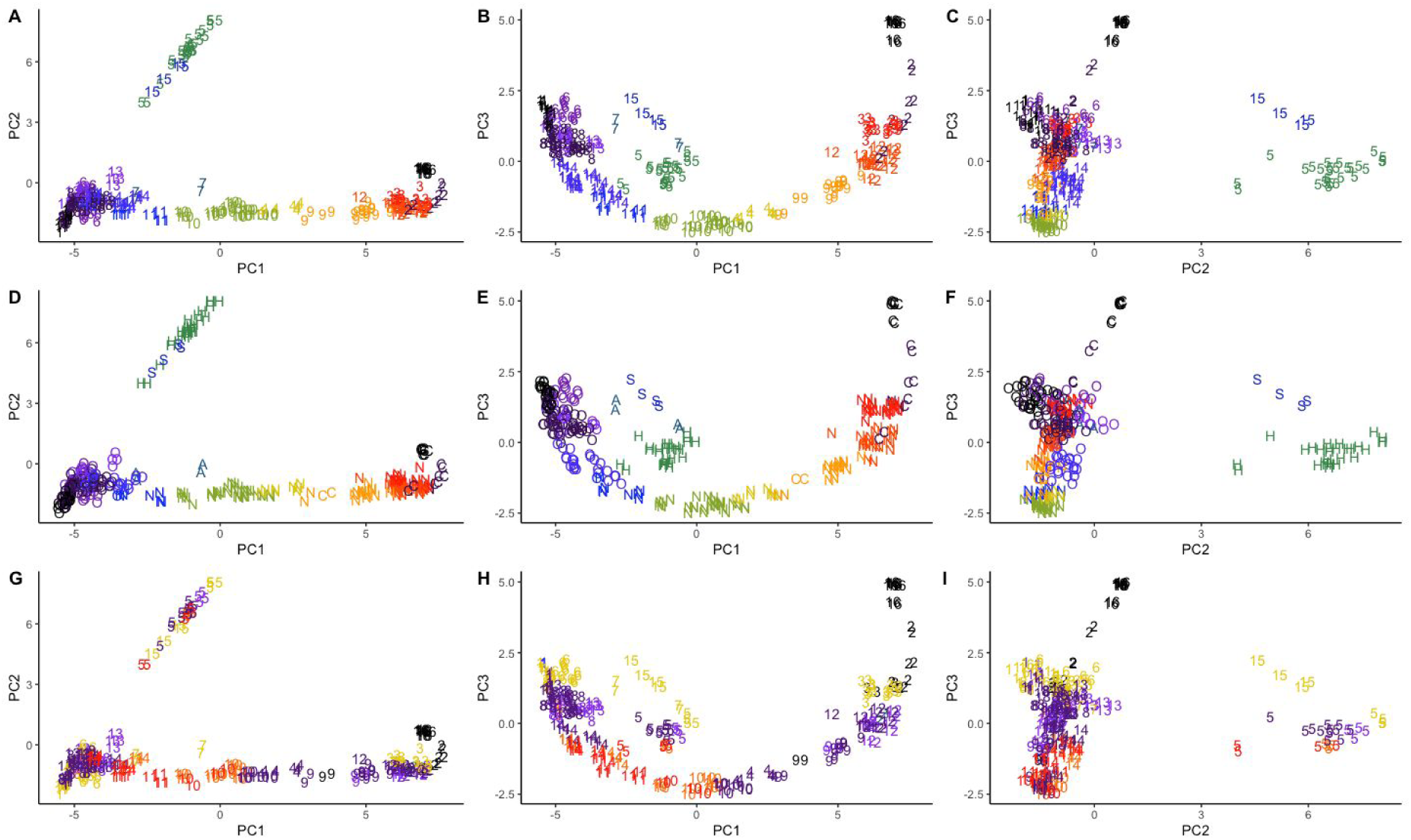
PCA analysis of 289 bulk RNA-seq samples. **A-C).** Labels in the plot correspond to the tissue cluster number which each sample belongs to. D-F) Each sample was labeled according to the first letter of the tissue of origin, which are; **s**keletal muscle, **h**eart, **a**drenal gland, **n**euronal (forebrain, hindbrain, midbrain, neural tube), **c**ortex and **o**thers. G-I). Each sample was labeled according to the a sample batch ID number, showing that the samples do not cluster by batch group.

**Figure S4.**
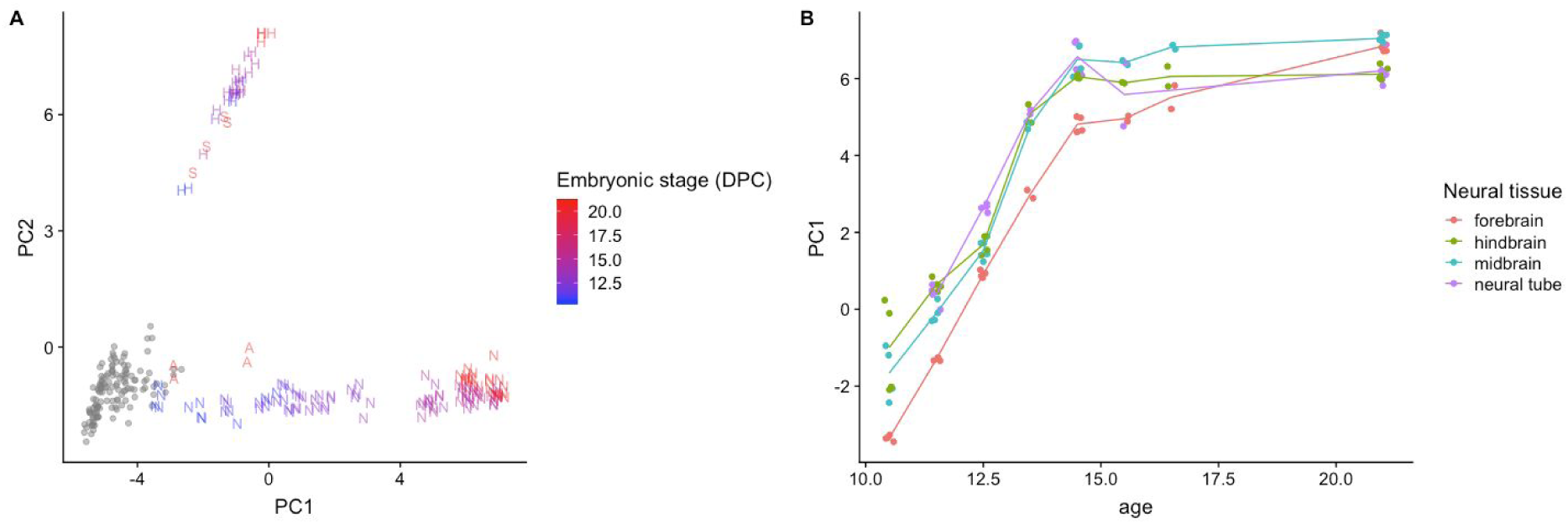
A) PCA plot where only the ENCODE samples are shown. Neuronal samples are color coded on a blue to red scale based on developmental time. B) Relationship between PC1 and mouse developmental stage (age) of ENCODE samples.

**Figure S5.**
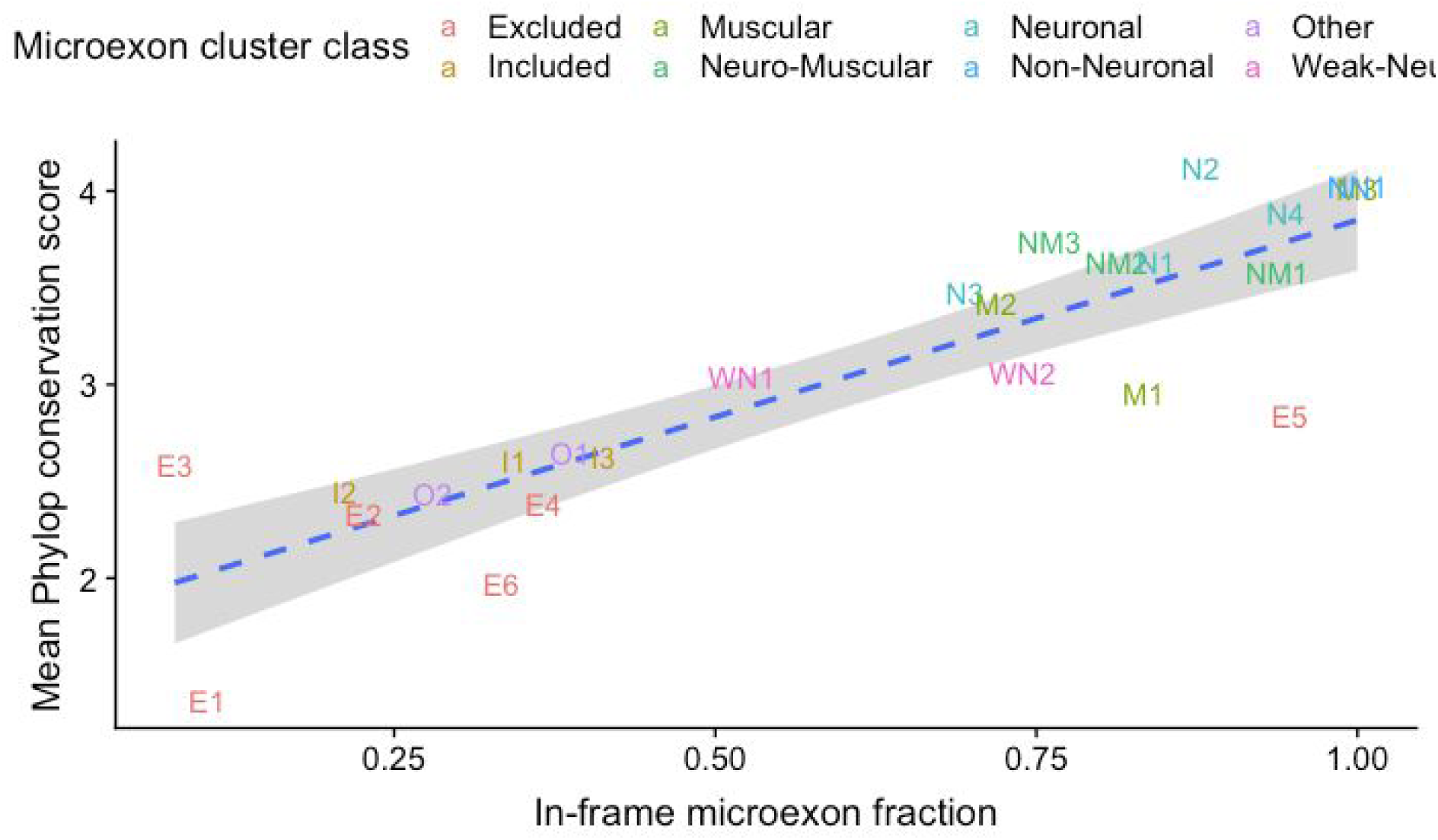
Relationship between mean conservation score (PhyloP) and fraction of in-frame microexons for different microexon clusters and developmental stages.

**Figure S6.**
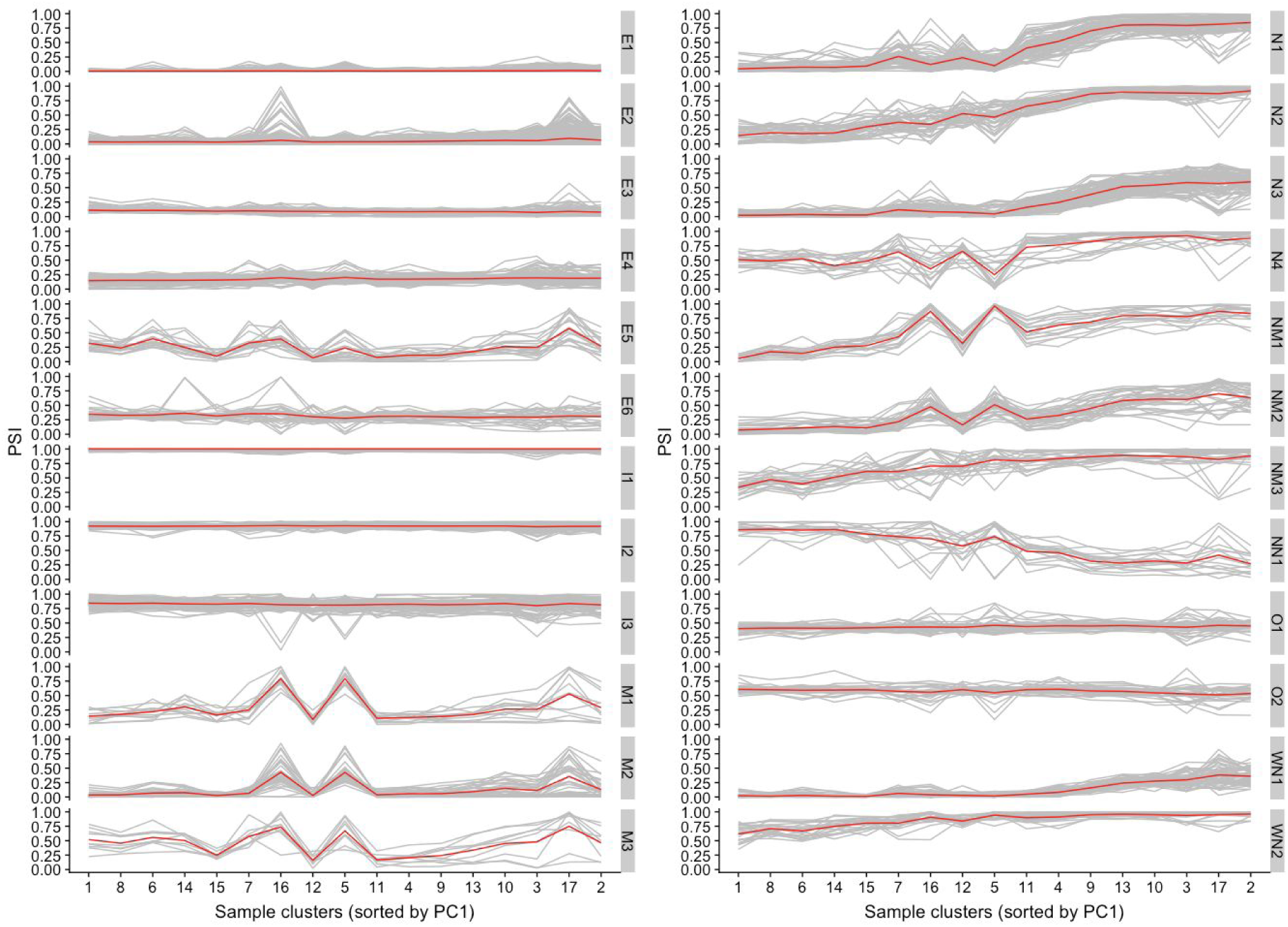
Average Microexon PSI for each microexon cluster across the different sample clusters are shown in red. Grey lines show the average PSI of individual microexons.

**Figure S7.**
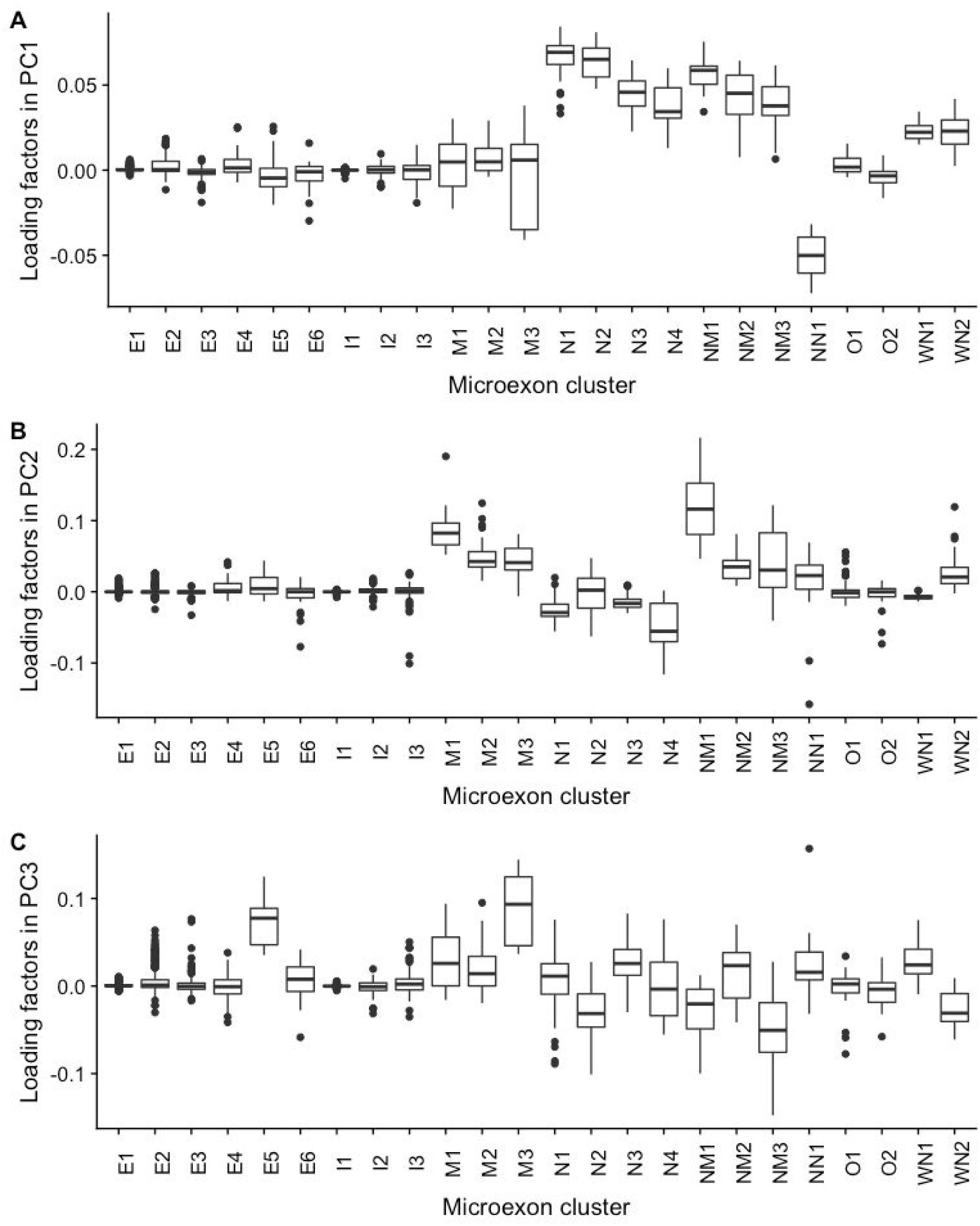
Boxplots showing PC1-3 loading factors for the different microexon clusters.

**Figure S8:**
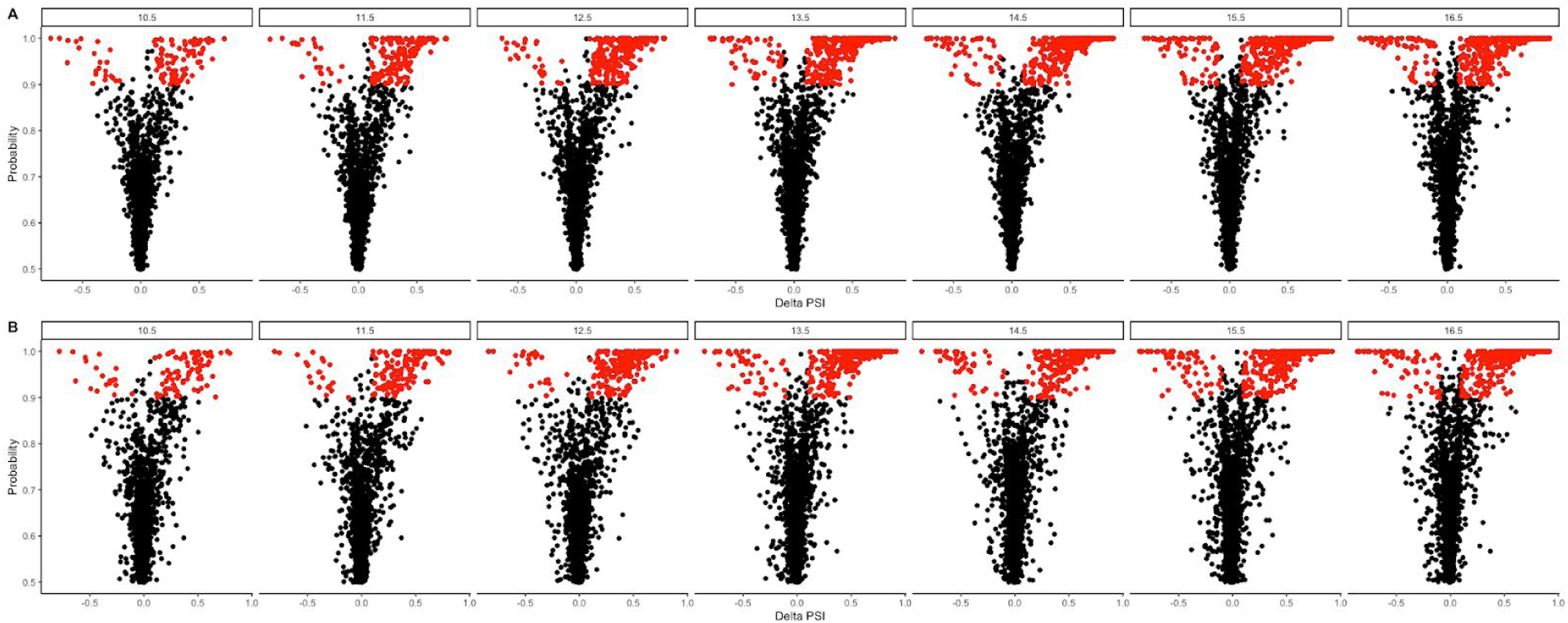
Volcano plots displaying microexon delta PSI values between neuronal samples of different developmental stages and control sample group, and their estimated probability to be differentially included. A) Whipped PSI estimations B) MicroExonator PSI estimations.

**Figure S9:**
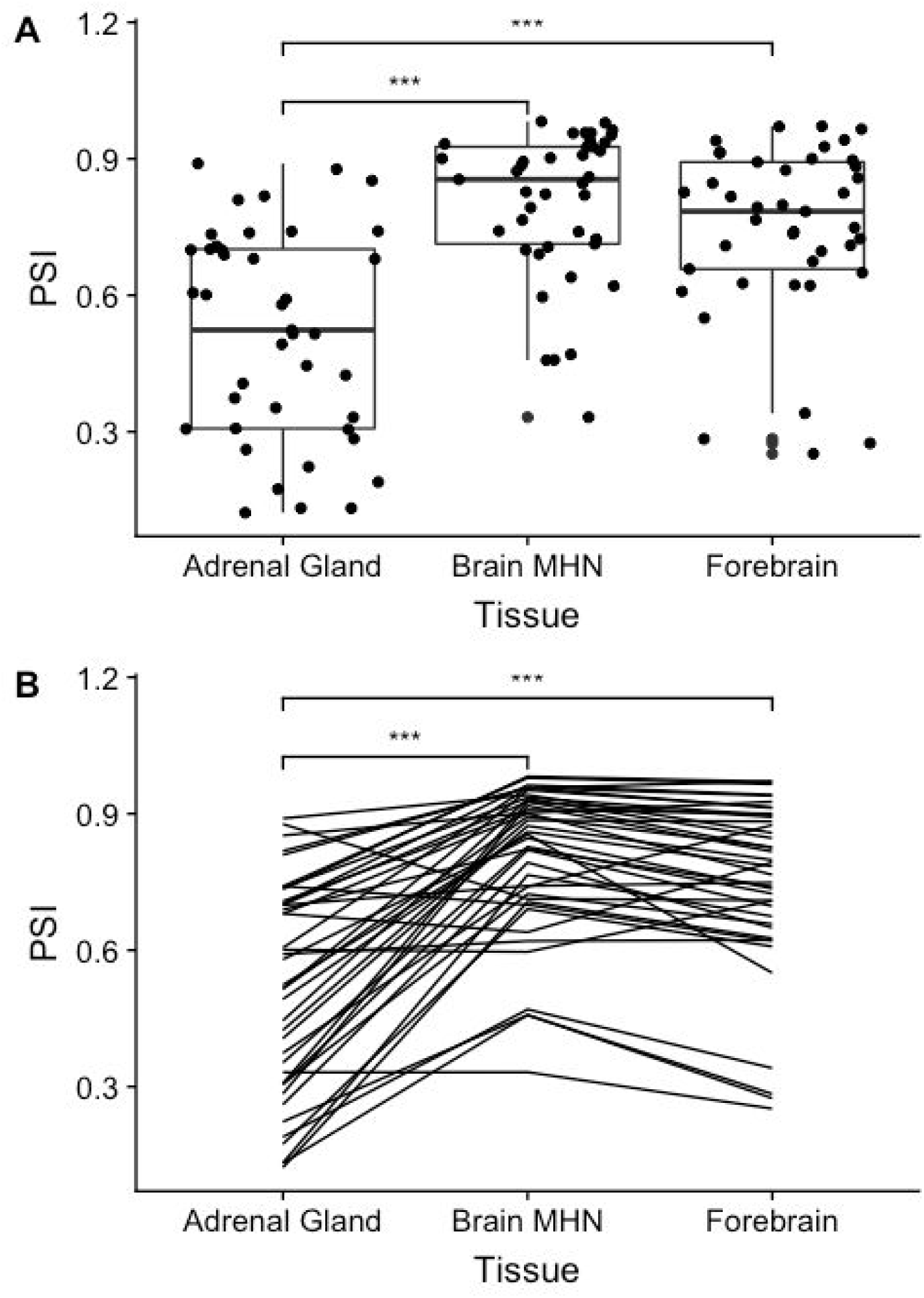
Differences in PSI values between adrenal gland, brain MHN and forebrain tissues. A) Boxplots showing PSI values of each group. Dots represent mean PSI values of each differentially included microexon across the different groups. B) Line plot showing the average PSI variation of each differentially included microexon across the different sample groups. Significant p-values are denoted by * (>0.05), ** (>0.01) and *** (>0.001).

**Figure S10.**
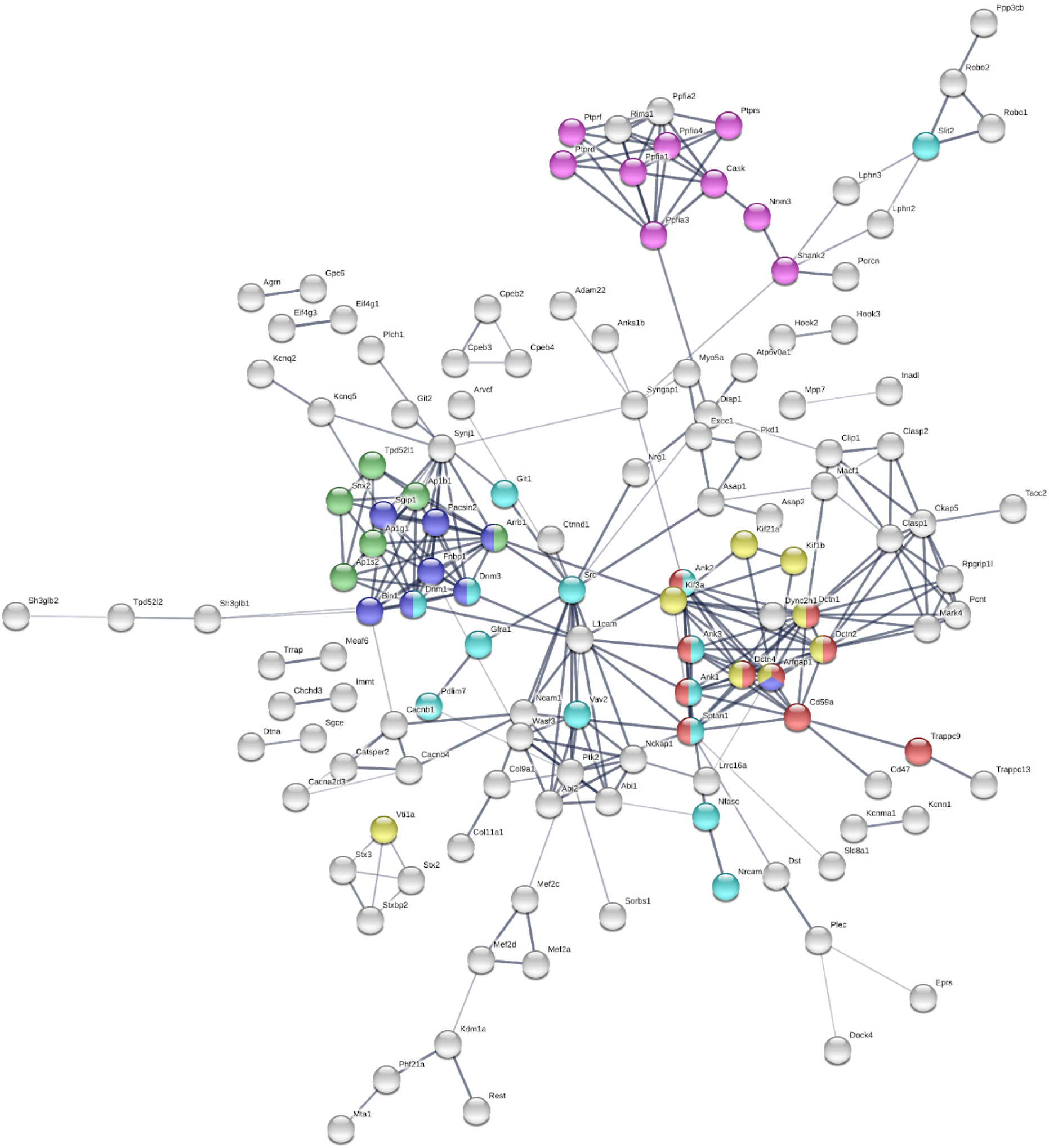
String PPI network of genes that were detected to have differentially included microexons between the control group and neuronal samples. Colors represent different Reactome pathways that were enriched on the network; Axon guidance (light blue), Protein-protein interactions at synapses (pink), ER to Golgi anterograde transport (red), Clathrin-mediated endocytosis (dark blue), Golgi associated vesicle biogenesis (green), Intra-Golgi and retrograde Golgi-to-ER traffic (yellow).

**Figure S11.**
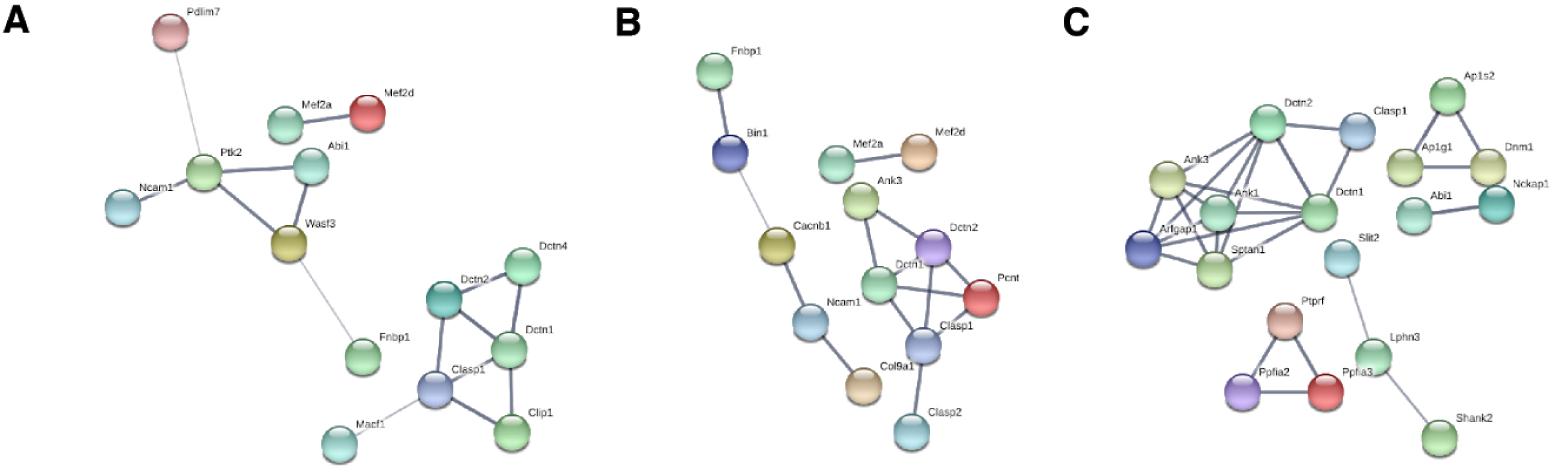
PPI network corresponding to the group of genes that were detected to have differentially included microexons between the control groups and A) Hearth B) Skeletal muscle C) Adrenal gland.

**Figure S12:**
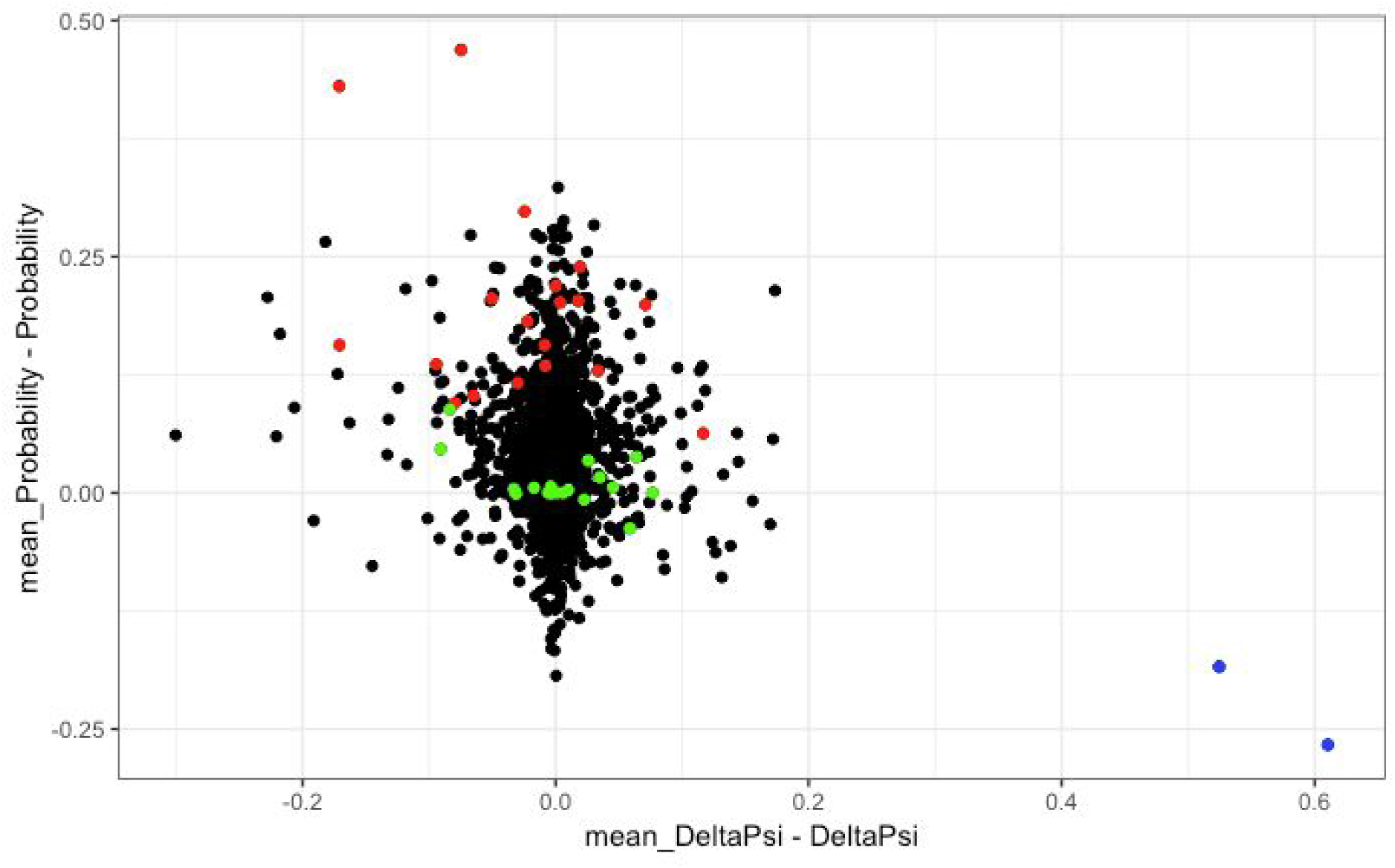
Scatter plot displaying the difference in probability score versus the difference in PSI. Red represents, green represents, blue represents, black represents no significant change.

**Table S1.**
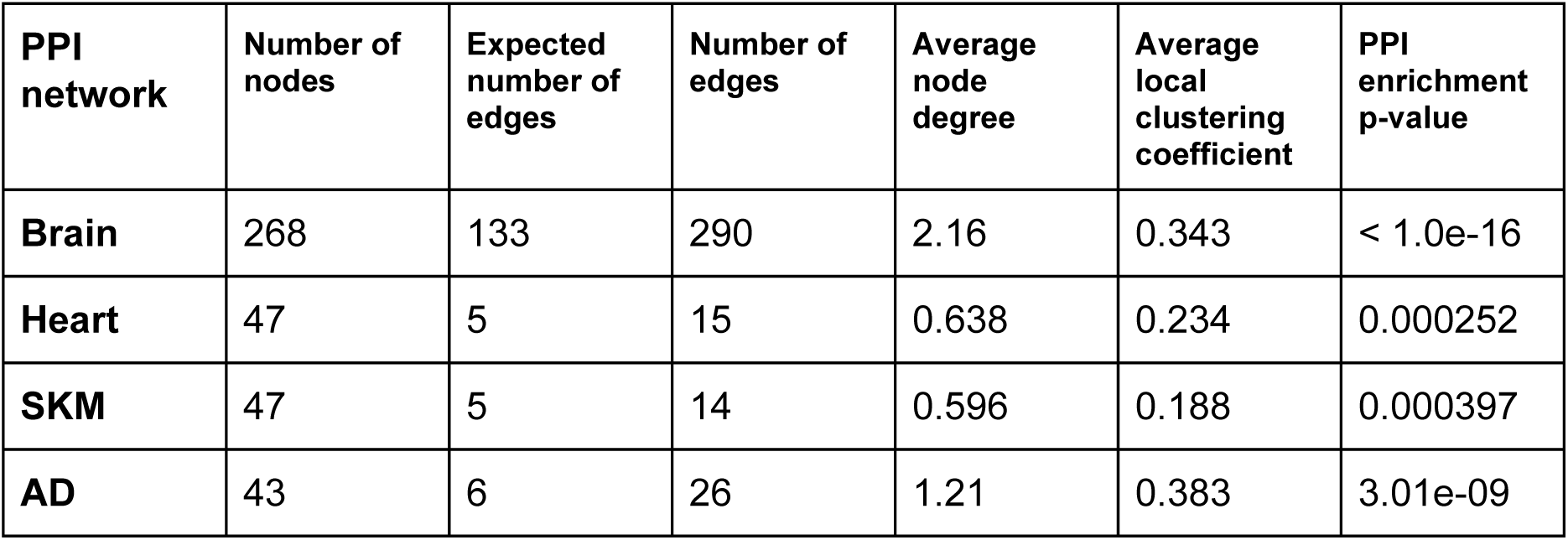
PPI network summary statistics reported by STRING.

